# Structure of the Visual Signaling Complex between Transducin and Phosphodiesterase 6

**DOI:** 10.1101/2020.04.01.020651

**Authors:** Yang Gao, Gözde Eskici, Sekar Ramachandran, Frédéric Poitevin, Alpay Burak Seven, Ouliana Panova, Georgios Skiniotis, Richard A. Cerione

**Author notes:** These authors contributed equally to this work (Yang Gao, Gözde Eskici, Sekar Ramachandran). Correspondence (R.A.C.), (G.S.).

## Abstract

Heterotrimeric G proteins communicate signals from activated G protein-coupled receptors to downstream effector proteins. In the phototransduction pathway responsible for vertebrate vision, the G protein-effector complex is comprised of the GTP-bound transducin α subunit (Gα_T_·GTP) and the cyclic GMP (cGMP) phosphodiesterase 6 (PDE6), which stimulates cGMP hydrolysis to transmit signals to the optic nerve. Here we report a cryo-electron microscopy (cryoEM) structure of PDE6 complexed to GTP-bound Gα_T_. The structure reveals two Gα_T_·GTP subunits engaging the PDE6 hetero-tetramer at both the PDE6 catalytic core and the PDEγ subunits, driving extensive rearrangements to relieve all inhibitory constraints on enzyme catalysis. Analysis of the conformational ensemble in the cryoEM data highlights the dynamic nature of the contacts between the two Gα_T_·GTP subunits and PDE6 that support an alternating-site catalytic mechanism.

## INTRODUCTION

Heterotrimeric G proteins, composed of a guanine-nucleotide-binding alpha subunit (Gα) and the constitutively associated beta and gamma subunits (Gβγ), function as cellular transducers that relay signals from a wide range of extracellular hormonal and sensory stimuli detected by G protein-coupled receptors (GPCRs) (Oldham and Hamm, 2008). Activated GPCRs catalyze the exchange of bound GDP for GTP on the Gα subunit, leading to the dissociation of the heterotrimer. The resulting GTP-bound Gα subunit (Gα·GTP), and in some cases the Gβγ subunit complex, subsequently engage a range of downstream effector proteins to trigger an array of intracellular signaling responses. Based on primary sequence similarity of the Gα subunits, heterotrimeric G proteins are divided into 4 classes: G_S_, G_i/o_, G_q/11_ and G_12/13_ (Simon et al., 1991). Several recent high-resolution structures of various GPCRs in complex with different G proteins have yielded important information on how this class of receptors selectively couple to and activate their signaling partners (Gao et al., 2019; García-Nafría and Tate, 2019; Kato et al., 2019; Krishna Kumar et al., 2019; Maeda et al., 2019). By contrast, structural information about how activated Gα subunits engage and activate full-length downstream effectors are limited to a crystal structure of Gα_q_ in a complex with phospholipase C-β3 (PLCβ3) (Lyon et al., 2013) and a recent cryoEM structure of a Gα_S_-adenylyl cyclase 9 (AC9) complex (Qi et al., 2019). Here, we present a cryoEM structure of a signaling complex between transducin, a G_i/o_ family member, and its full-length effector enzyme, cGMP phosphodiesterase 6 (PDE6). The findings elucidate an essential step in the signaling mechanism responsible for vertebrate vision and enhance our understanding of the structural basis of G protein-mediated signal transduction.

## RESULTS AND DISCUSSION

### Isolation and structure determination of the Gα_T_·GTP-PDE6 complex

In retinal rods (Figure 1A), the absorption of a photon by the GPCR rhodopsin (Rho) enables the receptor to bind and activate its cognate G protein signaling partner transducin (G_Τ_), thereby promoting the exchange of GDP for GTP within its α subunit (Gα_T_) (Stryer, 1991). GTP-bound Gα_T_ (Gα_T_·GTP) then activates PDE6 by relieving the inhibitory constraints imposed by the two identical PDEγ subunits (PDEγ, 10 kD) on the non-identical catalytic α and β subunits (PDEα and PDEβ, 99 kD and 98 kD, respectively). The stimulation of PDE6 enzymatic activity (i.e. the hydrolysis of cGMP to GMP) by Gα_T_·GTP reduces the local cytosolic cGMP concentration, leading to closure of cGMP-gated cation channels on the plasma membrane and hyperpolarization of rod photoreceptor cells to initiate the signal sent to the optic nerve.

**Figure 1.**
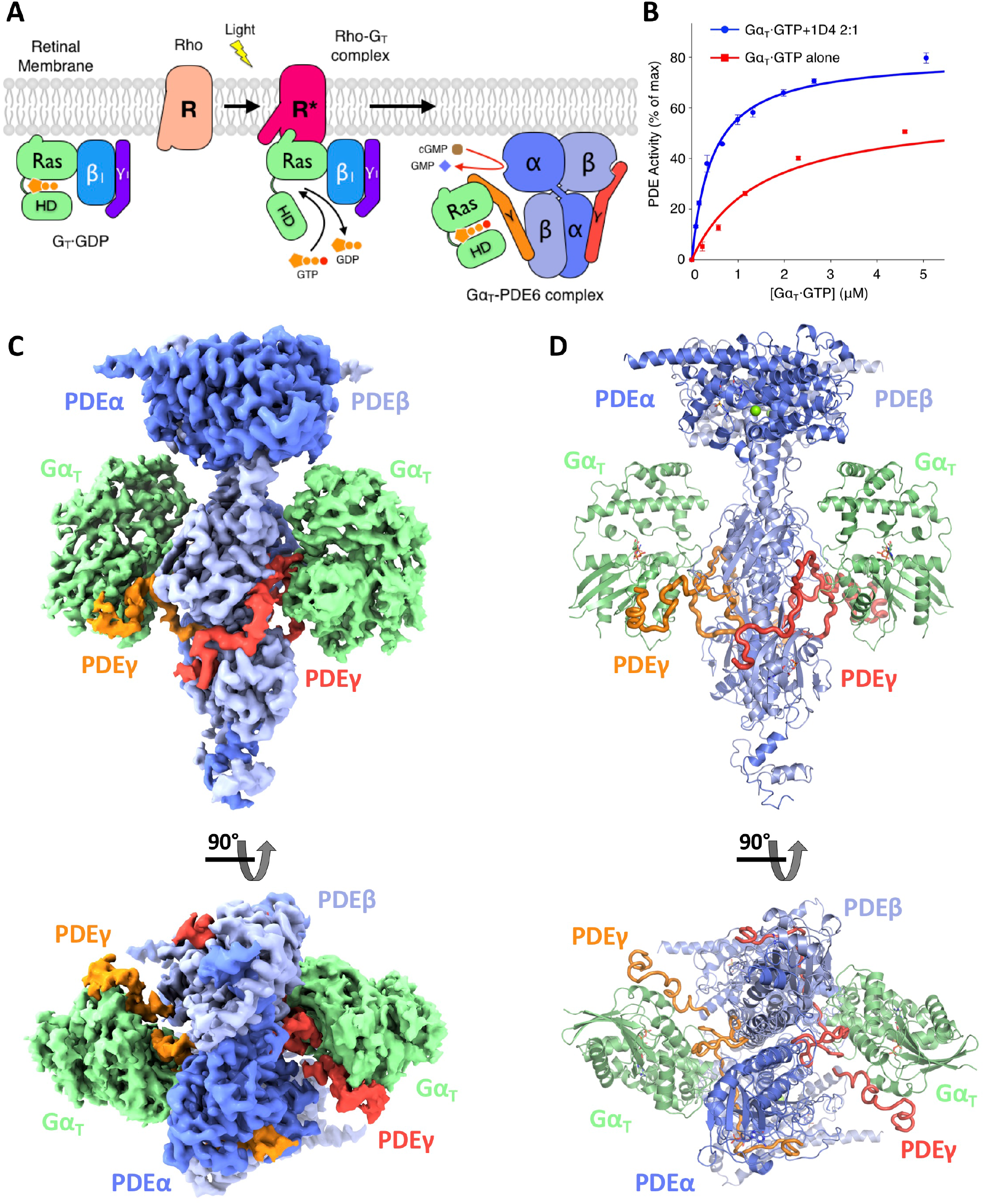
CryoEM structure of the transducin (Gα_T_)-phosphodiesterase 6 (PDE6) complex. (**A**) Schematic illustration of the vertebrate visual phototransduction pathway. Dark-adapted and light activated rhodopsin (Rho) are depicted as R (pink) and R* (red). Also depicted are the Ras and helical domains of the Gα_T_ subunit (green), the Gβ_1_ (cyan) and Gγ_1_ (purple) subunits of transducin, as well as the PDEγ subunits (orange and red), and the PDEα (dark blue) and PDEβ (light blue) catalytic and regulatory domains of PDE6. (**B**) PDE6 activity assays with the addition of varying concentrations of either Gα_T_·GTP or the Gα_T_·GTP-1D4 2:1 complex. (**C** and **D**) Orthogonal views of the cryoEM density map (C) and structural model (D) of the Gα_T_-GTP-PDE6 complex colored by subunit (Gα_T_·GTP subunits in green, PDEα in dark blue, PDEβ in light blue, PDEγ in orange and red).

To form a complex between Gα_T_·GTP and PDE6, we utilized native bovine PDE6 purified directly from bovine retina and a recombinant Gα_T_ subunit that contains 18 corresponding residues from Gα_i1_ (Figure S1A), allowing for its expression in *E. coli*. (Milano et al., 2018; Skiba et al., 1996). Two GTP hydrolysis-defective mutations (R174C and Q200L) were also introduced to maintain Gα_T_ in an activated GTP-bound state (Majumdar et al., 2006). The GTP-bound recombinant Gα_T_ subunit can activate PDE6 as effectively as GTPγS-bound native bovine retinal Gα_T_ (Figure S1B). Previous biochemical studies have suggested that the heterodimeric PDE6 has two Gα_T_·GTP binding sites with different binding affinities (Clerc et al., 1992). Of note, binding of two Gα_T_·GTP subunits are required to achieve maximal Gα_T_-stimulated PDE6 activity (Qureshi et al., 2018). However, maximal activation of the native enzyme by Gα_T_·GTP is consistently 50% of the activity measured when PDE6 is treated with trypsin, whereby both inhibitory PDEγ subunits are removed by protease-digestion (Wensel and Stryer, 1986). These observations suggest that the two catalytic domains of PDE6 are not simultaneously activated by two GTP-bound Gα_T_ subunits, pointing to an alternating-site catalytic mechanism. Interestingly, we previously showed that when an antibody recognizing the C-terminus of Gα_T_ was incubated at a 1:2 ratio with Gα_T_·GTPγS, it resulted in a two-fold enhancement of Gα_T_·GTPγS-stimulated PDE6 activity, matching the levels achieved when PDEγ subunits are removed through trypsinization (Phillips et al., 1989). These results suggested that the bivalent antibody, by presenting two Gα_T_·GTPγS subunits to PDE6, was able to relieve the inhibitory constraints imposed by both of the PDEγ subunits on the holo-enzyme.

Our early attempts to reconstitute a stable and homogeneous Gα_T_·GTP-PDE6 complex with a 2:1 stoichiometry for structural studies were unsuccessful, presumably because Gα_T_·GTP subunits dissociate from activated PDE6 during size exclusion chromatography as they alternate between higher and lower affinity states. Inspired by our earlier observations, we hypothesized that antibody presentation of two Gα_T_ subunits may stabilize a complex. To this end, we utilized a commercially available antibody (1D4) that recognizes the C-terminal 9-amino-acid sequence of rhodopsin (Molday and Molday, 2014), and attached the 1D4 epitope to the C-terminus of recombinant Gα_T_·GTP with a linker that afforded flexible tethering of Gα_T_ subunits (Figure S1A). This strategy allowed us to recapitulate the antibody potentiation and achieve a full activation of PDE6, with the complex formed between one 1D4 antibody and two Gα_T_·GTP subunits (Figure S1D). Importantly, this system significantly increased the affinity between Gα_T_·GTP and PDE6 as determined in PDE6 activity assays (Figure 1B). We were thus able to form a stable Gα_T_·GTP-PDE6 complex (Figures S1C and S1E), using native bovine PDE6 and the epitope-tagged recombinant Gα_T_·GTP in the presence of the 1D4 antibody and the PDE5/6 orthosteric inhibitor vardenafil, which occupies the substrate-binding sites on the PDEα and PDEβ subunits. 2D classification and averaging of cryoEM projections revealed well defined structural features for the complex, with the exception of the 1D4 antibody, which was highly flexible and was averaged out (Figure S2A). A 3D reconstruction using 143,125 particle projections yielded a final map of the Gα_T_·GTP-PDE6 complex with an indicated global resolution of 3.2 Å (Figure 1C and Figures S2B to S2D), facilitating the refinement of a near atomic resolution model (Figure 1D and Figures S3A to S3C).

### Structural basis of PDEγ extraction by Gα_T_·GTP

The structure of the Gα_T_·GTP-PDE6 complex reveals two Gα_T_·GTP subunits that are simultaneously bound to either side of the PDE6 hetero-tetramer with a pseudo two-fold symmetry (Figure 1D). The Gα_T_ subunits form extensive interactions with the inhibitory PDEγ subunits, burying interface areas of 1346 Å^2^ and 1385 Å^2^, respectively, on each side. Additionally, Gα_T_ forms limited contacts with the large catalytic PDEα and PDEβ subunits, with interface areas of 143 Å^2^ and 159 Å^2^, respectively.

The most striking feature of the Gα_T_·GTP-PDE6 complex involves the orientation of the Gα_T_·GTP subunits on PDE6 relative to the membrane plane. PDE6 is prenylated at the C-termini of the PDEα and PDEβ subunits and is peripherally attached to the rod photoreceptor outer segment membranes (Figure 2A). Compared to the recent cryoEM structure of the rhodopsin-transducin complex (Gao et al., 2019), and based on the presumed position of the membrane bilayer, each Gα_T_·GTP subunit in the Gα_T_·GTP-PDE6 complex rotates approximately 150° away from the conformation of Gα_T_ when bound to the photoreceptor (Figures 2A and 2B). The nearly upside-down orientation of Gα_T_·GTP in the Gα_T_·GTP-PDE6 complex allows the Ras domain of Gα_T_ to displace the C-terminal end of the PDEγ subunit away from its position that blocks substrate (cGMP) access to the catalytic sites on the PDEα and PDEβ subunits, resulting in a striking 60 Å movement of the PDEγ C-terminus when compared to a recent cryoEM structure of the inactive PDE6 enzyme (Figure 3A). The retinal Gα_T_, though N-terminally modified by either myristoylation or by one of three other less hydrophobic fatty acids (Neubert et al., 1992), associates very weakly with the membrane and easily dissociates from rod outer segment membranes when in the GTP-bound state. The recent Gα_S_-AC9 complex structure (Qi et al., 2019) shows that the Gα_S_ subunit is also detached from the membrane (Figure 2C). Thus, a relatively short-lived displacement from the membrane surface may be a common feature for enabling GTP-bound Gα subunits to engage and regulate the activities of their effector proteins.

**Figure 2.**
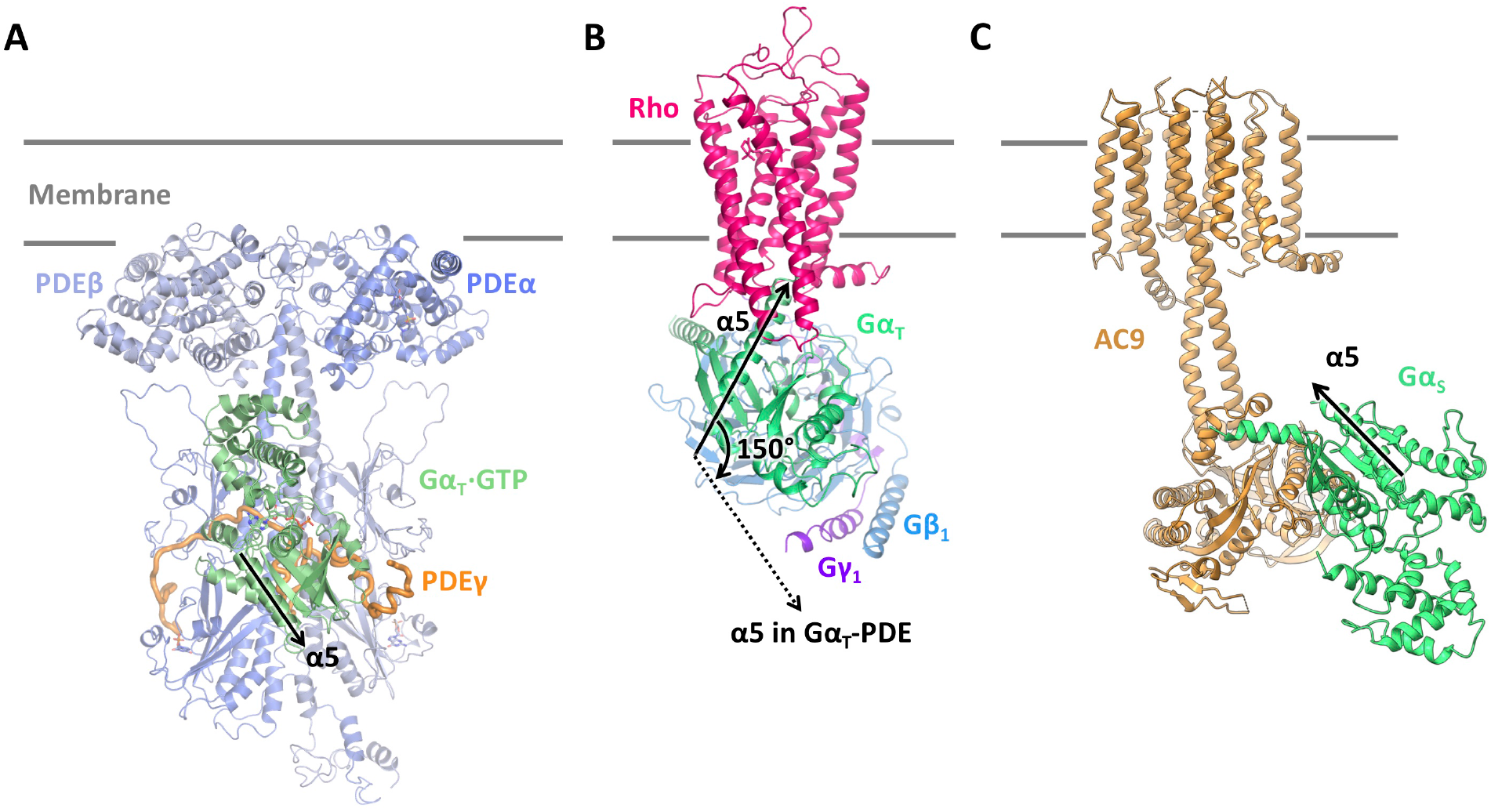
Comparison the orientation of Gα subunit in the Gα_Τ_·GTP-PDE6 complex with that in other complexes. (**A**) Orientation of the Gα_T_ subunit in the Gα_Τ_·GTP-PDE6 complex with the α5 helix conformation indicated by an arrow. (**B**) Orientation of the Gα_T_ subunit in the rhodopsin (Rho)-G_T_ complex (PDB: 6OYA). The orientation of the α5 helix in the Rho-G_T_ complex (solid arrow) is compared with that in the Gα_Τ_·GTP-PDE6 complex (dashed arrow). (**C**) Orientation of the Gα_S_ subunit in the Gα_S_-AC9 complex (PDB: 6R3Q) with the α5 helix conformation indicated by an arrow.

**Figure 3.**
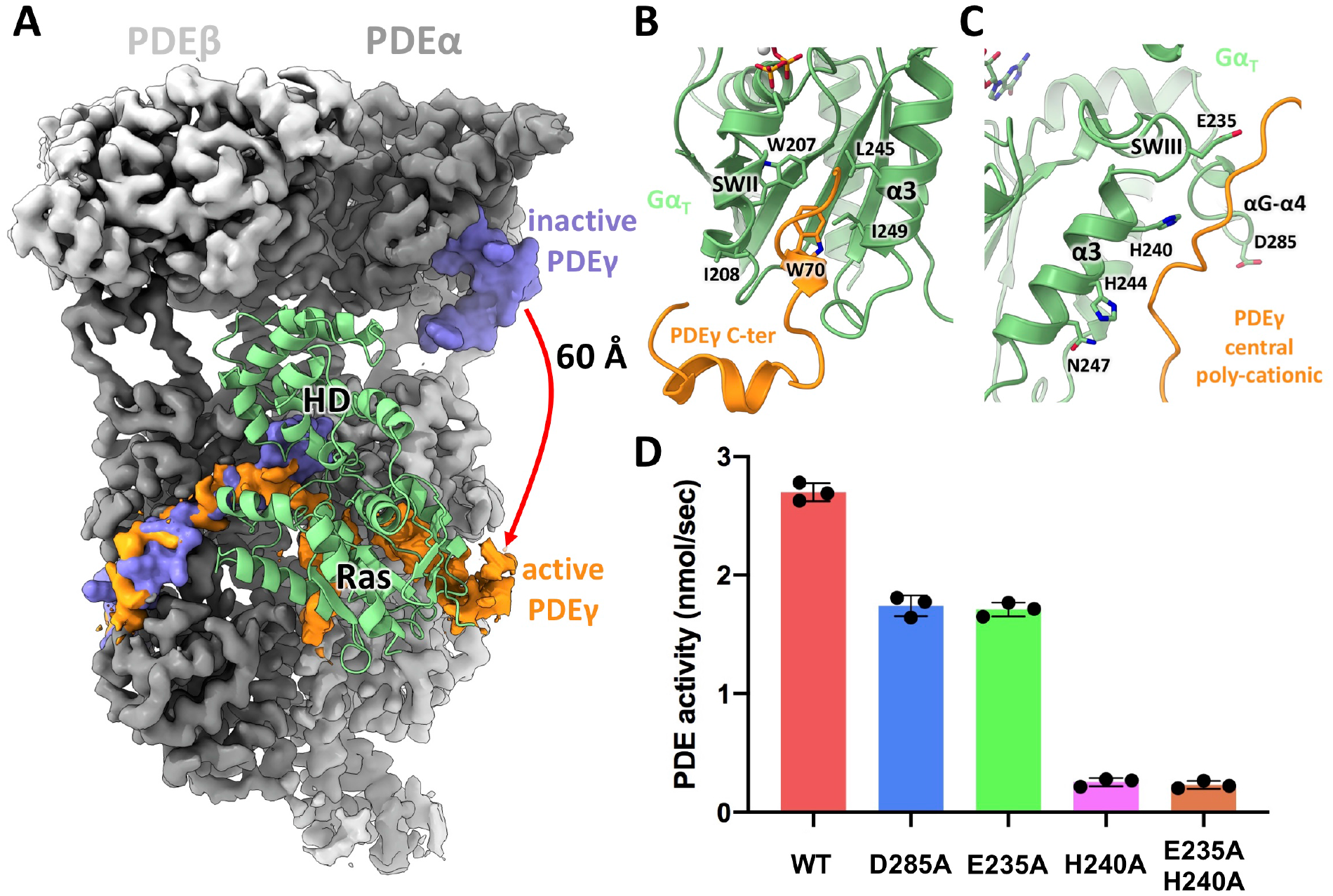
Interactions between the Gα_T_ Ras domain and PDE6. (**A**) Displacement of the C-terminus of PDEγ upon Gα_T_ binding (PDEα and PDEβ are shown as dark and light grey densities respectively, the Gα_T_·GTP subunit is shown as green ribbons, and PDEγ subunits in the inactive (PDB: 6MZB) and activated states are shown as blue and orange densities respectively. (**B**) Interactions between the PDEγ C-terminus and the Switch II and α3 helix regions of Gα_T_. (**C**) Key residues from the α3 helix, Switch III and αG-α4 loop of Gα_T_ help stabilize PDEγ. (**D**) Mutating PDE-interacting residues from the Gα_T_ Ras domain significantly reduces Gα_T_-induced PDE6 activity.

The interactions between the Gα_T_ Ras domain and the C-terminus of PDEγ involve the key Trp70 residue from PDEγ inserting into a hydrophobic pocket formed by W207 and I208 from the Switch II (SWII) region, and L245 and I249 from the α3 helix of Gα_T_·GTP (Figure 3B). This is similar to what was observed in a previous X-ray crystal structure of GDP·AlF_4_^−^-bound Gα_T/i1_ (a transition state mimic for GTP hydrolysis) in complex with the C-terminal peptide of PDEγ and the limit domain of the regulator of G protein signaling 9 (RGS9) protein (Gα_T/i1_-PDEγ C-ter-RGS9) (Slep et al., 2001). The upside-down conformation of Gα_T_·GTP in the Gα_T_·GTP-PDE6 complex allows for additional interactions between the Ras domain and the central polycationic region of PDEγ (residues 24-46). This region, which has been shown biochemically to contribute to PDE6 inhibition and interactions with activated Gα_T_ (Artemyev and Hamm, 1992), is not resolved in the inactive PDE6 structure and its binding site on Gα_T_ has remained unknown. In our structure, the unique conformation of Gα_T_·GTP complexed to PDE6 positions residues H240, H244 and N247 of the α3 helix, E235 of the switch III region (SWIII), and D285 of the αG-α4 loop, to form potential polar interactions with the polycationic regions of PDEγ (Figure 3C). When the Gα_T/i1_-PDEγ C-ter-RGS9 structure is aligned onto the inactive PDE6 structure, there is no overlap between the surfaces on PDEγ involved in binding Gα_T_ and its interactions with the catalytic PDEα/β subunits (Figure S4A). Moreover, the residues positioned to contact the PDEγ polycationic region are located on the side of the Gα subunit facing away from this region (Figure S4B). Therefore, the interactions between the Gα_T_ Ras domain and the PDEγ polycationic region observed in the Gα_T_·GTP-PDE6 complex structure are only possible when Gα_T_·GTP adopts the upside-down conformation. The structure suggests that these additional interactions act as a leveraging point to pull Gα_T_·GTP together with the PDEγ C-terminus away from the catalytic domains of PDE6.

In support of the role of this interface, mutating either E235 of SWIII or D285 of the αG-α4 loop to alanine (Ala) reduced the ability of Gα_T_·GTP to stimulate PDE6 activity; while mutation of the Gα_T_ α3 helix residue H240 to Ala almost completely abolished Gα_T_·GTP-stimulated activity (Figure 3D). In addition, mutating H244 and N247 within the α3 helix to their corresponding residues in Gα_i1_ (H244K and N247D) was reported to significantly impair the ability of Gα_T_ to activate PDE6 (Natochin et al., 1998). Both the SWIII region and the αG-α4 loop were previously shown using a Gα_S_/Gα_i2_ chimera to be essential for adenylyl cyclase activation (Berlot and Bourne, 1992), and a F312Y mutation in the αG-α4 loop of Gα_S_, which replaces the phenylalanine with the corresponding tyrosine residue in Gα_i1_, markedly compromised the ability of Gα_S_ to stimulate adenylyl cyclase (Milano et al., 2018). These results indicate that the SWIII region and the αG-α4 loop represent a common interface for interactions between GTP-bound Gα subunits and their downstream effectors. In the Gα_S_-AC9 complex structure, neither of these regions is in close proximity to the Gα_S_-AC9 interface (Figure S4C), suggesting that there might be additional intermediate states along the activation pathway of adenylyl cyclase that would allow for the interactions between these regions in Gα_S_ and its effector. Taken together, the interactions between the PDEγ polycationic region and the Gα_T_ Ras domain observed in the Gα_T_·GTP-PDE6 complex appear to be essential for Gα_T_·GTP-stimulated activation of PDE6.

In the inactive PDE6 structure (Gulati et al., 2019), the PDEγ subunit spans the entire length of each catalytic subunit (Figure S5A). The C-terminus of each PDEγ subunit binds at the entrance of the active site of each catalytic domain, while the N-terminus of each PDEγ interacts with the GAFa domain, forming a lid that covers the allosteric cGMP-binding site (Figure S5B). In the Gα_T_·GTP-PDE6 complex, the N-terminus of each PDEγ subunit forms similar interactions with each GAFa domain that has an allosteric site occupied by a cGMP molecule. However, removal of the PDEγ C-terminus from the catalytic domain relieves the constraints imposed on the catalytic subunits, resulting in an apparent 4.3 Å elongation of the PDEα/β heterodimer in the Gα_T_·GTP-PDE6 complex when compared to the inactive PDE6 structure (Figure S5A). The tandem GAF domains have previously been suggested to play an inhibitory role in regulating the activity of the catalytic domains (Zhang et al., 2008). Elongation of the catalytic subunits would likely weaken the interactions between the GAF and catalytic domains, thereby helping to promote PDE6 activation.

### Vardenafil binding in PDE6

Within the catalytic domains, we observe clear densities for the therapeutic molecule vardenafil (Figure 4A). Inhibitors such as sildenafil and vardenafil have been widely used in treatment of pulmonary hypertension and erectile dysfunction by targeting PDE5, which is involved in modulating smooth muscle contraction (Maurice et al., 2014). However, due to the high degree of sequence homology between the catalytic domains of PDE5 and PDE6, inhibitors that target PDE5 can also bind to PDE6 with high affinity (Tejada et al., 2001). As a result of this cross reactivity, common side effects of these inhibitors include blurred vision, color vision alteration and even damage to the optic nerve (Moschos and Nitoda, 2016). The major conformational difference between the catalytic domains in the Gα_T_·GTP-PDE6 complex structure and a previous crystal structure of the PDE5 catalytic domain complexed with vardenafil (Sung et al., 2003) lie in the H-loop, a variable region that flanks the active site (Figure 4B). In PDE6, the H-loop adopts an open conformation, whereas in PDE5 the position of the H-loop shifts by 20 Å toward the active site. To avoid steric clash, the sulfonamide group at the 5’ position of vardenafil’s phenyl ring in PDE5 is rotated by nearly 180° and packs against the α10 helix in the catalytic domain (Figure 4B). This observation points to the possibility of designing better selective PDE5 inhibitors with less PDE6 cross reactivity by increasing the rigidity of the sulfonamide group of vardenafil to favor its PDE5-interacting conformation.

**Figure 4.**
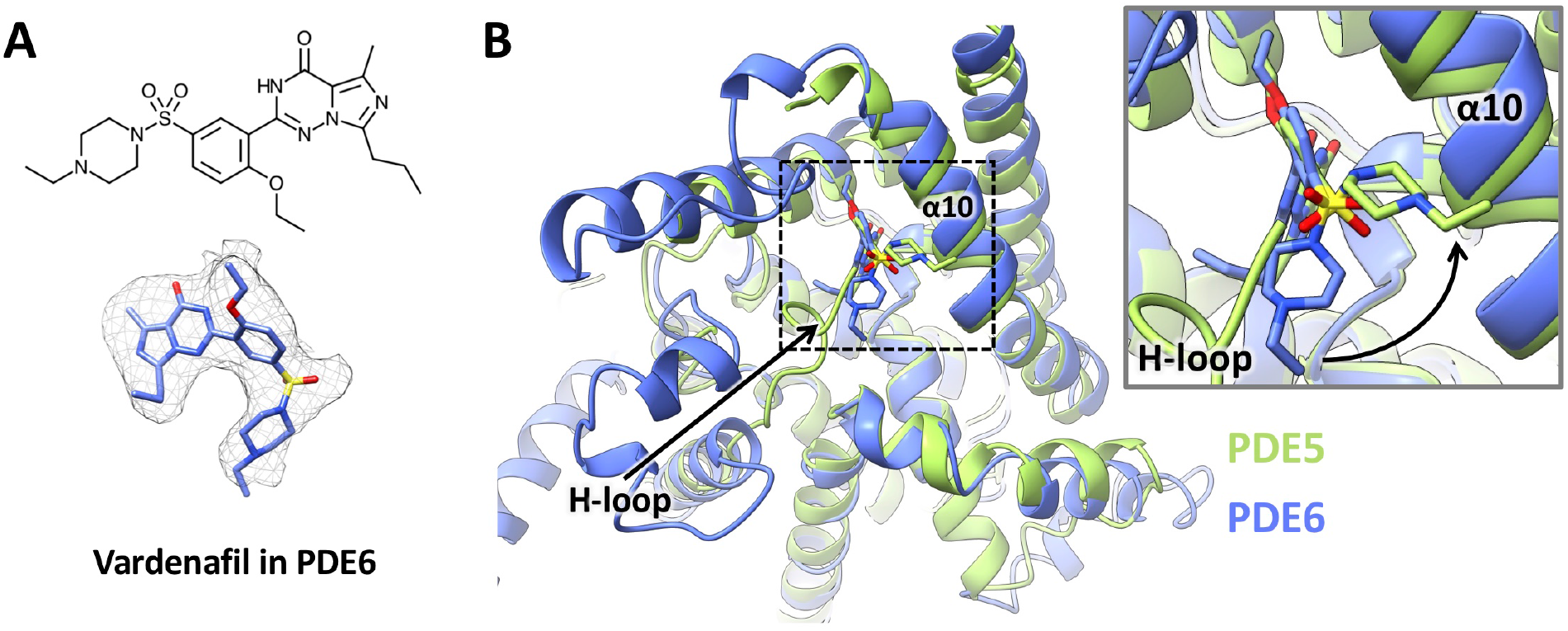
Binding of vardenafil in the Gα_T_·GTP-PDE6 complex. (**A**) Model and isolated density map of vardenafil in the Gα_T_·GTP-PDE complex. (**B**) Comparison between the catalytic domain from the Gα_T_·GTP-PDE6 complex structure (colored blue) and the crystal structure of the PDE5 catalytic domain complexed with vardenafil (colored green, PDB: 1UHO).

### Gα_T_·GTP αHD and PDE6 interactions bridge two PDE6 active sites and promote activation

In addition to forming extensive contacts with the PDEγ subunits, Gα_T_·GTP also interacts with the catalytic PDEα and PDEβ subunits via the α helical domain (αHD). Alignment of the Ras domains from the crystal structure of Gα_T_·GTPγS (Noel et al., 1993), with the Gα_T_·GTP structure in the PDE6 complex (Figure 5A), reveals that the αHD in Gα_T_·GTP extends slightly upward to form interactions with the N-terminal end of the α6 helix in the adjacent PDE catalytic subunit. Thus, upon displacing PDEγ away from the PDEα subunit, the αHD of a Gα_T_·GTP subunit would be positioned to interact with the catalytic domain of PDEβ. This upward extension places polar residues from the αB helix of the αHD, such as D93, S94, Q97 and D98, within interaction proximity to residues R652 and R653 from PDEα, and residues R650 and R651 from PDEβ. Mutating these αHD residues to Ala causes a modest reduction in the Gα_T_·GTP-stimulated PDE6 activity (Figure 5B). A previous study examining the heterologous expression of the Gα_T_ αHD moiety showed that the αHD synergistically enhances Gα_T_·GTPγS activation of PDE6 through interactions with the PDE6 catalytic core (Liu and Northup, 1998). These results suggest that the interactions between the αHD and PDE catalytic subunits, though relatively weak as observed in the cryoEM map for the Gα_T_·GTP-PDE6 complex, further contribute to PDE6 activation by Gα_T_·GTP. In the structure for inactive PDE6 (Gulati et al., 2019), the C-terminal ends of the α6 helix within the catalytic domains are directly involved in interactions with the C-termini of the PDEγ subunits (Figure S5C). Therefore, perturbation of the N-terminal end of the α6 helix by the αHD of a bound Gα_T_·GTP subunit could help weaken the interaction between PDEγ and the catalytic domain on the C-terminal end of the α6 helix, thereby promoting the binding of the other Gα_T_·GTP subunit to that PDEγ subunit and its displacement from the catalytic domain.

**Figure 5.**
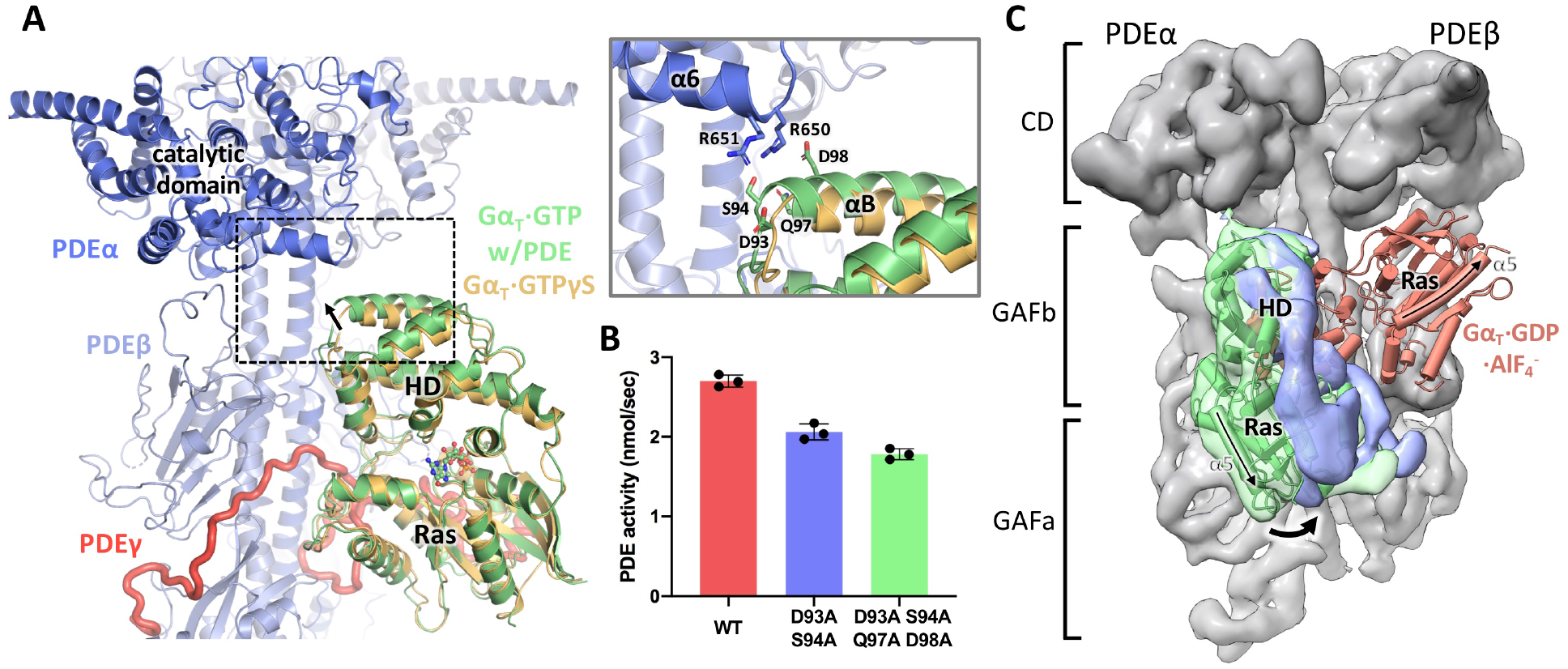
Interactions between the Gα_T_ helical domain and PDE6. (**A**) Interactions between the Gα_T_ helical domain and the catalytic domain of PDE6 (PDEα in dark blue, PDEβ in light blue, PDEγ in red, and Gα_T_·GTP in green). The Gα_T_·GTPγS crystal structure (in yellow, PDB: 1TND) is overlaid on top of Gα_T_·GTP based on alignment of the Gα_T_ Ras domain). (**B**) Mutating helical domain residues that are contacting PDE impairs Gα_T_-induced PDE6 activity. (**C**) The range of motions of Gα_T_ in the complex revealed by multi-body refinement. The two extreme conformations of Gα_T_·GTP are shown as green and blue densities with the Gα_T_·GTP structure docked into the green density. The encounter conformation of Gα_T_ on PDE6 is shown as a pink cartoon using the Gα_T_·GDP·AlF_4_^−^ structure from the Gα_T_·GDP·AlF_4_^−^-PDEγ-C-ter-RGS9 domain complex (PDB: 1FQJ), superimposed onto the inactive PDE6 structure (PDB:6MZB), based on alignment of the PDEγ C-ter peptide.

### Motions of the Gα_T_·GTP subunits bound to PDE6 shed light on alternating-site catalysis

During processing of cryoEM projections for 3D reconstructions, we observed significant flexibility in the Gα_T_·GTP subunits bound to the PDE6 complex (Figure S2A). This is also reflected in local resolution measurements of our final 3D map, which display higher resolution at the core of the PDEα and PDEβ subunits and relatively lower resolution at the outer edges of the two Gα_T_ subunits (Figure S2D). To better understand the extent of the movements of the Gα_T_·GTP subunits in the complex with PDE6, we employed multi-body refinement (Nakane et al., 2018) to visualize their motions (Figure S6). Specifically, the Gα_T_·GTP-PDE6 complex was divided into three bodies, namely the PDEα/β subunits in complex with the N-termini of two PDEγ subunits, and the two Gα_T_·GTP subunits in complex with the C-terminal and central polycationic regions of PDEγ (Figure S6B). We then analyzed the results through principal component (PC) analysis as implemented in Relion (Nakane et al., 2018), followed by an additional independent component (IC) analysis (Hyvarinen, 1999) in order to probe the existence of concerted motions of the Gα_T_·GTP subunits bound to PDE6 (Figures S6A to S6E, see Methods section for a detailed account). This analysis shows that the Gα_T_·GTP subunits undergo an alternating translational motion on the surface of PDE6, such that when one Gα_T_·GTP subunit undergoes movement the other remains fixed in one position (Figure S6B, Movies S1 and S2). This interpretation of the relative motions of the Gα_T_·GTP subunits is further supported by 3D maps that were refined from equally sized subsets of particles clustered along each IC (Figure S6F and S6G). The 3D maps show that the motions in the first two ICs correspond to the two Gα_T_·GTP subunits following pendulum-like movements by pivoting around the αHD-PDE6 catalytic domain interface (Figure 5C) in alternating fashion. The observed movements of the Gα_T_·GTP subunits further support an alternating-site catalysis mechanism within the two active sites of PDE6. The initial concept for such a mechanism stems from the finding that protease digestion of the PDEγ subunits consistently yields twice the catalytic activity observed upon stimulation by Gα_T_·GTP, and from the ability of a bivalent antibody to simultaneously present two Gα_T_·GTP subunits to PDE6 and thus give rise to a 2-fold enhancement in catalytic activity. Moreover, previous biochemical studies have suggested that binding of the first Gα_T_ subunit only provides limited stimulatory activity on PDE6, with the simultaneous binding of two Gα_T_·GTP subunits being necessary to achieve maximal Gα_T_-stimulated PDE6 activity (Clerc et al., 1992; Qureshi et al., 2018).

A Gα_T_ peptide encompassing the α4 helix (residues 293-314) has been reported previously to stimulate PDE6 activity (Rarick et al., 1992). The downward rotation of the Gα_T_·GTP subunits that we observe in the Gα_T_·GTP-PDE6 complex would likely bring this region in contact with the PDE6 GAFb domain, which modulates the activity of the catalytic domain of the opposite PDE6 catalytic subunit. Thus, the potential contacts between the Gα_T_·GTP subunits and the GAF domains, as well as interactions between the Gα_T_ αHD and the PDE6 catalytic domain, would provide the structural basis for the crosstalk between the two PDE6 catalytic subunits (Figure 6). Our results suggest that binding of the first Gα_T_·GTP on one side of PDE6, not only removes PDEγ from blocking the active site of the corresponding catalytic domain, but also primes the other catalytic domain for activation by engaging it through the αHD to relieve the inhibition of the GAF domains. Subsequently, the second Gα_T_·GTP would only need to displace the C-terminal end of PDEγ from the other subunit to initiate catalysis. As the interactions between Gα_T_·GTP and regions of PDE6 other than the PDEγ C-terminus are flexible, it is likely that in the absence of an antibody holding both Gα_T_·GTP subunits in place, the two Gα_T_ subunits would alternatively induce the activating conformation observed in the Gα_T_·GTP-PDE6 complex structure.

**Figure 6.**
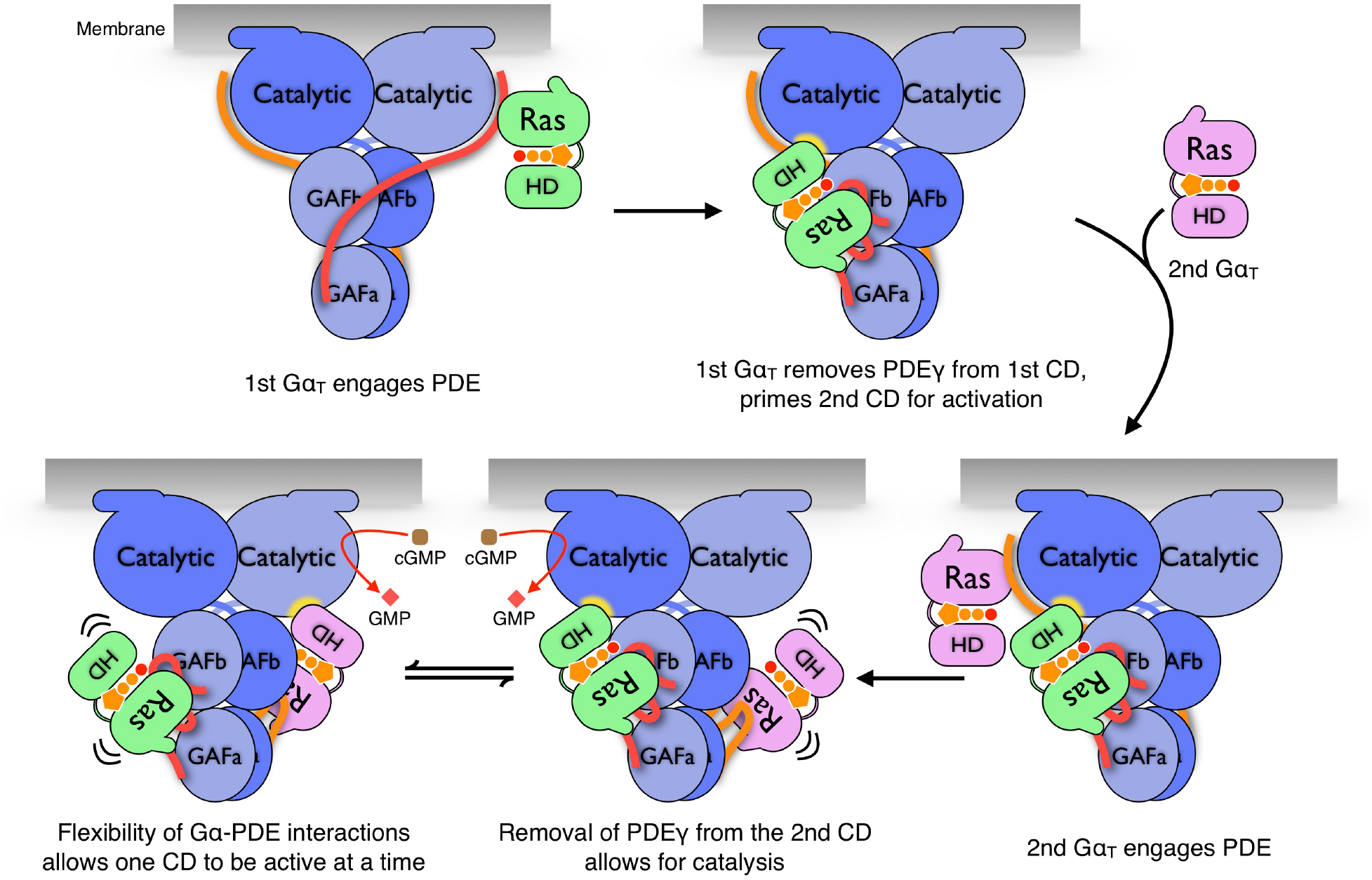
Schematic illustrating the mechanism of Gα_T_-induced activation of PDE6. The Ras and helical domains of the two bound Gα_T_·GTP subunits are depicted in green and blue. The catalytic and GAF domains of PDEα and PDEβ are shown in dark and light blue, respectively, and the PDEγ subunits are in red and orange. The first GTP-bound Gα_T_ engages PDE6 at the C-terminus of one PDEγ, removes the PDEγ peptide from the corresponding PDE6 catalytic domain (CD) and primes the second CD for activation. The second GTP-bound Gα_T_ extracts PDEγ from the second CD allowing for cGMP hydrolysis at this site. The interactions between Gα_T_ and PDE catalytic subunits are flexible (shown by the squiggly lines), and in the absence of the 1D4 antibody, only one CD is fully active at a time.

## Conclusions

The Gα_T_·GTP-PDE6 cryoEM structure provides a starting mechanistic framework for a key event in visual phototransduction. The structure and its underlying dynamics represent a snapshot into the concerted interactions between an activated G protein and its full-length phosphodiesterase effector protein. The upside-down orientation adopted by the two Gα_T_·GTP subunits in the Gα_T_-PDE6 complex not only allows for the engagement between the Ras domain of a Gα_T_ subunit and the C-terminal and central polycationic regions of a PDEγ subunit, but also brings the αHD of Gα_T_ in position to interact with the catalytic domain on the opposite side of the PDE6 complex. The requirement for two Gα_T_ subunits to stimulate catalysis at each active site on PDE6 in an alternating manner would be advantageous in filtering out the noise from low-level spontaneous activation of Gα_T_ molecules. This mechanism would allow for the essential fast signaling turn-off required for visual phototransduction, whereby the RGS9 complex (He et al., 1998) needs to deactivate only one GTP-bound Gα_T_ molecule in order to attain a significant decrease in PDE6 activity. Given that our cryoEM data analysis shows that the coupling between Gα_T_·GTP and PDE6 is highly dynamic with pendulum-like motions, an upward rotation could potentially bring a Gα_T_·GTP subunit together with the C-terminus of a PDEγ subunit back to the active site of a catalytic subunit of PDE6 (Figure 5C). A recent mass-spectrometry analysis of a crosslinked complex between the GTP hydrolysis-transition state mimic, Gα_T_·GDP·AlF_4_^−^, and PDE6 revealed a conformation reminiscent of an encounter complex where a Gα_T_ subunit initially engages PDE6 at the C-terminus of a PDEγ subunit (Irwin et al., 2019). The RGS9 complex involved in deactivating GTP-bound Gα_T_ is attached to the membrane by the RGS9 anchor protein (R9AP) (Hu and Wensel, 2002). Alignment of previous crystal structures of the RGS9 complex (Cheever et al., 2008) together with the Gα_T_-RGS9 domain complex (Slep et al., 2001) onto the inactive PDE6 structure (Gulati et al., 2019), shows that RGS9 can fit perfectly parallel to the membrane plane (Figure S7). We thus postulate that the RGS9 complex would act on Gα_T_·GTP when it swings back up toward the membrane. The Gα_T_·GTP-PDE6 complex structure presented here provides a platform upon which additional structures of the PDE6 complex, trapped in different signaling states with other key regulatory proteins, can be added toward achieving a comprehensive view of the dynamic nature of this remarkable sensory response system.

## METHOD DETAILS

### Purification of PDE6 and transducin from bovine retina

PDE6, the retinal Gα_T_ subunit, and the Gβ_1_γ_1_ subunit complex were purified from bovine retina essentially as described (Phillips et al., 1989). 300 Dark-adapted bovine retina (W.L. Lawson Co., Lincoln, NE) were exposed to light and subjected to sucrose gradient ultra-centrifugation to prepare purified rod outer segment (ROS) membranes. ROS membranes were washed (3X) with 50 mL isotonic buffer (10 mM HEPES pH 7.5, 100 mM NaCl, 1 mM DTT, 5 mM MgCl_2_, 0.1 mM EDTA) and PDE6 was extracted from ROS membranes with three washes of 50 mL hypotonic buffer (10 mM HEPES pH 7.5, 1 mM DTT, 0.1 mM EDTA). The membranes were then washed with 100 mL GTP buffer (hypotonic buffer+100 μM GTP) to release the retinal Gα_T_ subunits and the Gβ_1_γ_1_ subunit complex. The hypotonic washes containing PDE6 were subjected to anion exchange chromatography through a 1 mL HiTrap Q HP (GE Healthcare) column, using Buffer A (20 mM Tris pH 8.0, 5 mM MgCl_2_, 1 mM DTT) and Buffer B (Buffer A + 1 M NaCl) to form the gradient. The eluted PDE6 was further purified with size exclusion chromatography using a Superdex 200 10/300 GL column (GE Healthcare) equilibrated with a buffer containing 20 mM Tris pH 8.0, 5 mM MgCl_2_, 1 mM DTT. PDE6 was concentrated to ~10 μM, flash-frozen and stored at −80°C with the addition of 10% glycerol. The GTP wash containing Gα_T_ and Gβ_1_γ_1_ was loaded onto a 5 mL HiTrap Blue HP (GE Healthcare) column to separate Gα_T_ from Gβ_1_γ_1_. The fractions containing Gα_T_ or Gβ_1_γ_1_ were further purified separately by anion exchange chromatography through a 5 mL HiTrap Q HP (GE Healthcare) column, using Buffer A (20 mM HEPES pH 7.5, 5 mM MgCl_2_, 1 mM DTT, 10% glycerol) and Buffer B (Buffer A + 1 M NaCl) to form the gradient. Both retinal Gα_T_ and Gβ_1_γ_1_ were concentrated to ~20 μM, flash-frozen and stored at −80°C.

### Purification of recombinant 1D4-tagged Gα_T_ subunit

Recombinant Gα_T_, containing a C-terminal 3-Ala linker with a 1D4 epitope tag (TETSQVAPA) and GTP hydrolysis-defective substitutions (R174C and Q200L) (Figure S1A), was expressed in *E. coli* BL21(DE3) competent cells and purified as described previously (Majumdar et al., 2006). The proteins were purified with anion exchange chromatography through a 5 mL HiTrap Q HP (GE Healthcare) column, using Buffer A (20 mM HEPES pH 7.5, 5 mM MgCl_2_, 1 mM DTT, 10% glycerol) and Buffer B (Buffer A + 1 M NaCl) to form the gradient. The eluted fractions were concentrated to about 20 μM, flash-frozen and stored at −80°C.

### Gα_T_·GTP-PDE6 complex formation and purification

1D4-tagged recombinant Gα_T_·GTP was washed with buffer containing 20 mM Tris pH 8.0 and 5 mM MgCl_2_ on a 10kD MWCO concentrator to remove DTT and then mixed with the 1D4 antibody (University of British Columbia, CA) in a 7:1 molar ratio and incubated on ice for 30 min. The mixture was loaded onto a Superdex 200 10/300 GL column (GE Healthcare) equilibrated with buffer containing 20 mM Tris pH 8.0, and 5 mM MgCl_2_ to purify the 2:1 Gα_T_·GTP-1D4 complex. PDE6 was washed with buffer containing 20 mM Tris pH 8.0, and 5 mM MgCl_2_ on a 100kD MWCO concentrator to remove DTT and mixed with an equal molar amount of the purified 2:1 Gα_T_·GTP-1D4 complex together with 10 μM of vardenafil (Sigma-Aldrich). The mixture was incubated on ice for 30 min and purified by gel filtration chromatography using a Superdex 200 10/300 GL column (GE Healthcare) equilibrated with buffer containing 20 mM Tris pH 8.0, 5 mM MgCl_2_ and 1 μM vardenafil. Peak fractions were pooled and concentrated with a 100kD MWCO concentrator to ~10 mg/mL.

### Gα_T_ subunit mutations and PDE6 activity assays

Point mutations were introduced onto a wild-type recombinant Gα_T_ construct that lacked the C-terminal 1D4 epitope tag and the GTP hydrolysis-defective substitutions (R174C and Q200L). Both the wild-type and mutant proteins were expressed and purified as described for the 1D4-tagged protein. The concentrations of the mutants were normalized to that of the wild-type protein based on the amplitude of intrinsic tryptophan fluorescence changes that occur as a result of conformational changes in the switch II region of Gα_T_ upon the addition of aluminum fluoride (AlF_4_^−^) (Majumdar et al., 2006). Fluorescence measurements were carried out on a Varian eclipse spectrofluorometer (excitation: 280 nm; emission: 340 nm). Gα_T_ (500 nM) was mixed with 1 mL of buffer containing 20 mM HEPES pH 7.5, 5 mM MgCl_2_ and 100 mM NaCl at room temperature and the fluorescence emission was monitored in real-time with the addition of AlF_4_^−^ (i.e. that forms from a mixture of 5 mM NaF and 50 μM AlCl_3_).

cGMP hydrolysis by PDE6 was measured and analyzed as described previously (Majumdar et al., 2006). Typically, in 200 μL of assay buffer containing 10 mM Tris pH 8.0, 2 mM MgCl_2_, and 100 mM NaCl, the GTPγS-loaded wild-type or mutant Gα_T_ subunit (1 μM) was incubated with 50 nM PDE6. The pH (in mV) was monitored in real time, and upon achieving a stable baseline, 5 mM cGMP was added and the decrease in pH was recorded for 150 sec. The buffering capacity of the mixture was obtained by adding NaOH (400 nmol). The hydrolysis rate of cGMP (nmol/sec) was determined from the ratio of the initial slope of the pH record (mV/sec) and the buffering capacity of the assay buffer (mV/nmol).

Trypsinized PDE6 was prepared by treating purified PDE6 (5 μM) in assay buffer with TPCK-treated trypsin (55 μg/mL) at room temperature. At the end of the incubation period soybean trypsin inhibitor (600 μg/mL) was added to quench the proteolytic reaction. The activity of trypsinized PDE6 was determined as described above without the addition of the Gα_T_ subunit. The PDE6 activities in the presence of varying concentrations of retinal Gα_T_·GTPγS, recombinant Gα_T_·GTP or the 2:1 Gα_T_·GTP-1D4 complex were measured as described above and normalized using the activity of trypsinized PDE to percent of maximal activity.

### CryoEM data collection and processing

A 3.5 μL solution of the Gα_T_·GTP-PDE6 complex (2 mg/mL) supplemented with 0.05% (w/v) β-octyl glucoside (Anatrace) was applied to freshly glow-discharged gold holey carbon grids (Quantifoil, Au-R1.2/1.3) under 100% humidity. Excess sample was blotted away for 2 seconds at 20°C, and the grids were subsequently plunged-frozen using a Vitrobot Mark IV (Thermo Fisher Scientific). A total of 1083 movies were recorded on a Titan Krios electron microscope (Thermo Fisher Scientific - FEI) operating at 300 kV with a calibrated magnification of x29,000 and corresponding to a magnified pixel size of 0.8521 Å. Micrographs were recorded using a K3 direct electron camera (Gatan) with a dose rate of 12 electrons/Å^2^/s and defocus values ranging from - 1.5 μm to −2.5 μm. The total exposure time was 4 s and intermediate frames were recorded in 0.07s intervals, resulting in an accumulated dose of 48 electrons per Å^2^ and a total of 57 frames per micrograph. Dose fractionated image stacks were subjected to beam-induced motion correction and filtered according to the exposure dose using MotionCor2 (Zheng et al., 2017). The sum of each movie was applied to CTF parameters determination by Gctf (Zhang, 2016). Auto-picked 841,967 particle projections were extracted and subjected to several rounds of reference-free 2D classification using Relion 3 (Zivanov et al., 2018). An initial model was computed from 325,795 particles using the Stochastic Gradient Descent algorithm implemented in Relion 3, and this model was used as an initial reference for 3D classification. Several rounds of 3D classification were performed to remove particles populating poorly defined classes. Conformationally homogeneous groups accounting for 143,125 particles were subjected to 3D masked refinement followed by map sharpening implemented in Relion 3. The resulting map had an indicated global nominal resolution of 3.5 Å. By using Relion 3.1, estimated CTF parameters were refined, and per-particle reference-based beam induced motion correction was performed using Bayesian polishing. The final map has a global resolution of 3.2 Å. Reported resolution is based on the gold-standard Fourier shell correlation (FSC) using the 0.143 criterion. Local resolution was estimated using the Relion 3.1 implementation.

### Model building and refinement

The initial model for the Gα_T_·GTP-PDE6 complex was constructed as poly-Ala chains based on the cryoEM structure of inactive PDE6 (PDB:6MZB) (Gulati et al., 2019) and the crystal structure of GTPγS-bound Gα_T_ (PDB: 1TND) (Noel et al., 1993), and manually docked into cryoEM density using Chimera (Pettersen et al., 2004). The model was then subjected to iterative rounds of automated refinement using Phenix real space refine (Adams et al., 2010), and manual building in Coot (Emsley and Cowtan, 2004). Sequence assignment was guided by bulky amino acid residues, such as Phe, Tyr, Trp and Arg. The final model was subjected to global refinement and minimization in real space in Phenix. Validation was performed in MolProbity (Chen et al., 2010) and EMRinger (Barad et al., 2015). The final refinement statistics are provided in Table S1.

### Flexibility analysis

The three masks were manually built in Chimera (Pettersen et al., 2004) from the consensus map. The consensus map and three body masks are input to the multi-body refinement tool in Relion (Nakane et al., 2018), which refines 6 parameters (3 translations and 3 rotations) for each of the three bodies with the following parameters: initial angular sampling 0.2, initial offset range 3, an initial offset step 0.75. We note ****X**** the resulting *m-by-n* array whose *i^th^* column *X_i_* contains the *m=18* rigid-body parameters assigned to the particle image *i*. The dataset is made of *n* (*n* = 143,125) ≫ *m* particle images. The dimensionality of the data is further reduced using principal component analysis (PCA) that yields a latent space of dimension 3 then interpreted using independent component analysis (ICA).

#### Principal Component Analysis

The principal component analysis (PCA) of ***X*** is similar to its singular value decomposition (SVD) that amounts to factorizing it as a product of three matrices: ***X*** = ***U∑V***^***T***^, after centering. Unitary matrices *m-by-m **U*** (resp. *n-by-m **V***) contain the left (resp. right) singular vectors of ***X***, and the diagonal matrix **∑** contains their corresponding singular values, sorted in decreasing order. The *k^th^* entry of the *j^th^* left eigenvector, *U*_*j*_*(k)*, is the contribution of the *k*^*th*^ rigid-body parameter to the *j*^*th*^ principal component. The *k*^*th*^ entry of the *j*^*th*^ right eigenvector, *V_*j*_(k)*, is the coordinate of the *k^th^* image particle along the *j^th^* principal component. The variance associated with the *j*^*th*^ component is the square of the *j*^*th*^ diagonal entry of **∑**. SVD was carried out in RELION (Nakane et al., 2018), generating the following files: ‘analyse_eigenvectors.dat’ that stores the left singular eigenvectors, and ‘analyse_projections_along_eigenvectors_all_particles.txt’ that stores the right singular vectors scaled by their respective singular values. For purposes of visualization and further analysis, we first recombine ****X**** from those files and carry SVD using the numpy (van der Walt et al., 2011) and scikit-learn (Pedregosa et al.) python libraries. PCA is often used to separate out signal-bearing components from noisy components (Colwell et al., 2014). However, PCA does not guarantee the signals to be carried by individual components. Rather the signals are mixed and spread on the principal components. Therefore, we used the first three eigenvectors from PCA with the highest variance to filter the data and the associated eigenvalues and the corresponding reduced coordinates were kept when recombining the data. We used a two-step un-mixing approach. <u>1. Data filtering:</u> We note 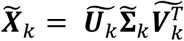, the filtered version of ***X*** where only the first *k* (*k* = 3) singular vectors and values are used. <u>2. Signal unmixing:</u> The resulting matrix was unmixed by Independent Component Analysis (ICA) (Hyvarinen, 1999) whose aim is to identify independent sources from multiple measurements of a mixture of signals.

#### Independent Component Analysis

Formally, ICA yields the *n-by-m **S*** source matrix after “whitening” (with a matrix ***W***) ***X*** and “un-mixing” it (with the transpose of a matrix ***M***): ***WX*** = ***MS***^***T***^. Whitening could be done in several ways, as long as the resulting covariance matrix is diagonal one. Un-mixing is achieved by maximizing the negentropy (i.e. reverse entropy) of each of the sources: sources that display a normal distribution have lowest negentropy. The negentropy of the *j*^*th*^ source can be defined as the Kullback-Leibler divergence between the density of the source ***S***_*j*_ and a normal equivalent ***N**j* (with same mean and variance as the source): *j*(**S***j*) = KL(p(**S**_*j*_)|p(**N**_*j*_)). ICA was performed on 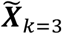 using the FastICA algorithm (Hyvarinen, 1999) available in scikit-learn (Pedregosa et al.). Formally, the vectors of the matrix **W**^−1^**M** are the ICA version of **U∑** in PCA and are referred to in this work as the independent components. Correspondingly, the vectors of ***S*** are the ICA equivalent of the vectors of ***V*** in PCA, and in this work are referred to as the coordinates of the image particles along the independent components. Three ICs were extracted, and they display a clear one-to-one mapping between each IC and the motion of each body (Figure S6B). The first two ICs have a large negentropy and correspond to each of the Gα_T_·GTP subunits experiencing an alternating translational motion on the surface of PDE6, such that one Gα_T_·GTP undergoes this movement at a given time while the other Gα_T_·GTP remains fixed (Figures S6B and S6C). The third IC, however, has negligible negentropy and corresponds to the rotation of PDE6 on itself (Figures S6B and S6C). As such it can be interpreted as a source of noise with large variance added to the signal carried by the first two ICs.

#### Local 3D reconstruction

It is expected that the proximity of particles in reduced coordinates reflects their similarity in conformation. Grouping particles based on their proximity, or clustering them, should thus allow to reconstruct 3D classes that reflect the true distribution of the imaged object in its conformational space. Here we binned the image particles based on the order of their coordinates along the first and then second independent components (IC). For each IC we defined 10 bins of roughly equal size (~10,000 particles per bin). The resulting particle set for each bin of each IC was given to RELION for 3D reconstruction.

**Figure S1.**
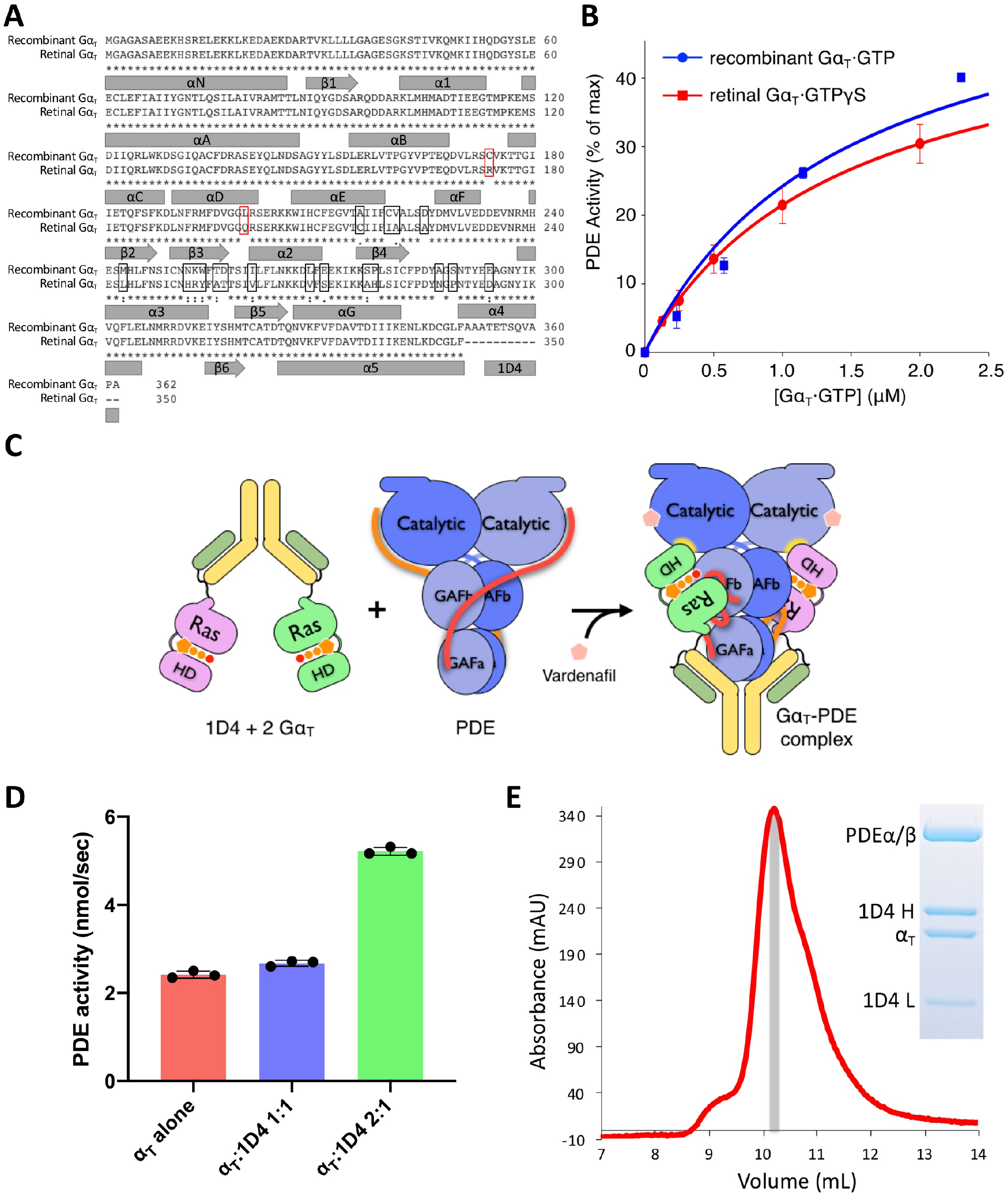
Purification of the Gα_T_·GTP-PDE6-1D4 complex, Related to Figure 1. (**A**) Protein sequence alignment of the 1D4-tagged recombinant Gα_T_ and bovine retinal Gα_T_. Rectangles (black outline) show the residues from Gα_i1_ introduced into the wild-type Gα_T_ backbone. The substitutions made to maintain recombinant Gα_T_ in an activated GTP-bound state are indicated by red rectangles. (**B**) PDE6 activity assays with either retinal Gα_T_ or the 1D4-tagged recombinant Gα_T_. (**C**) Schematic illustration of the purification of the Gα_T_·GTP-PDE6-1D4 antibody complex. (**D**) Gel filtration profiles and SDS-PAGE of the Gα_T_·GTP-PDE6-1D4 antibody complex.

**Figure S2.**
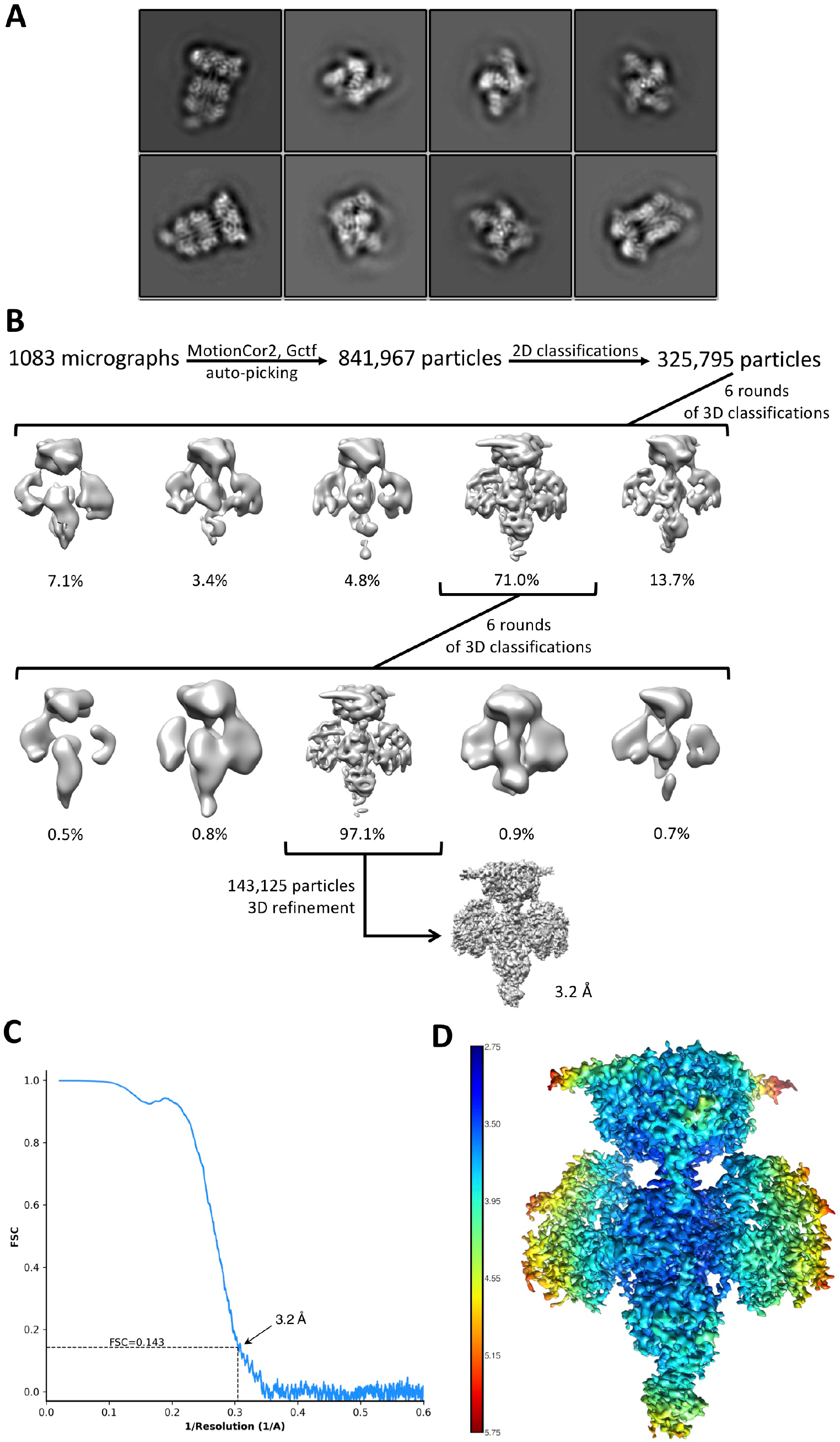
CryoEM of the Gα_T_·GTP-PDE6-1D4 complex, Related to Figure 1. (**A**) Representative 2D averages showing distinct secondary structure features from different views of the complex. (**B**) Flow chart of cryoEM data processing. (**C**) ‘Gold standard’ FSC curve indicating overall nominal resolutions of 3.2 Å using the FSC = 0.143 criterion. (**D**) Local resolution estimation of the cryoEM map.

**Figure S3.**
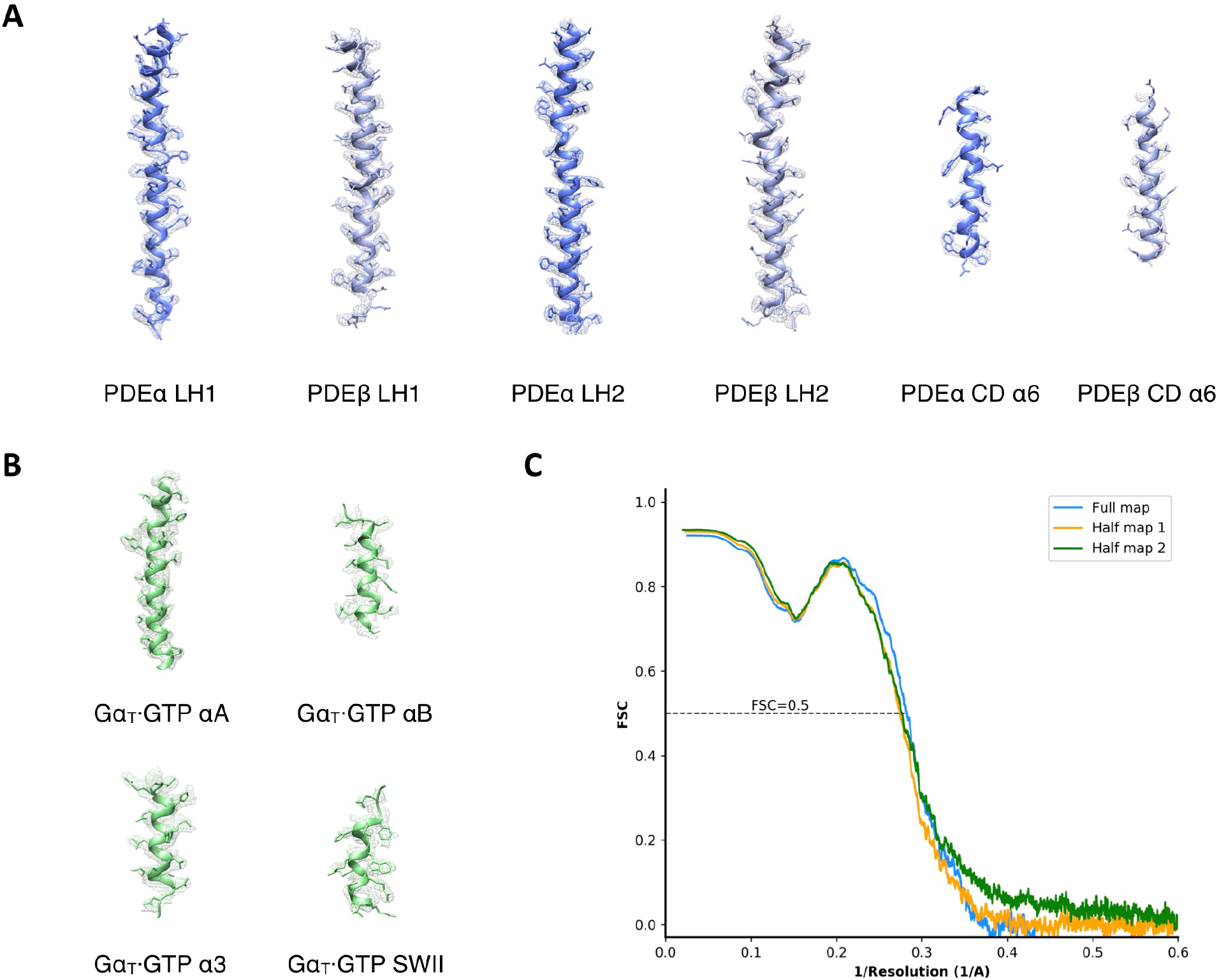
CryoEM map versus refined structure, Related to Figure 1. CryoEM densities and refined models for representative regions of PDE6 (**Α**) and Gα_T_·GTP (**Β**) in the complex. (**C**) FSC validation of the model versus cryoEM map, the model refined against the first half-map (work), and the latter model versus the second half-map (free).

**Figure S4.**
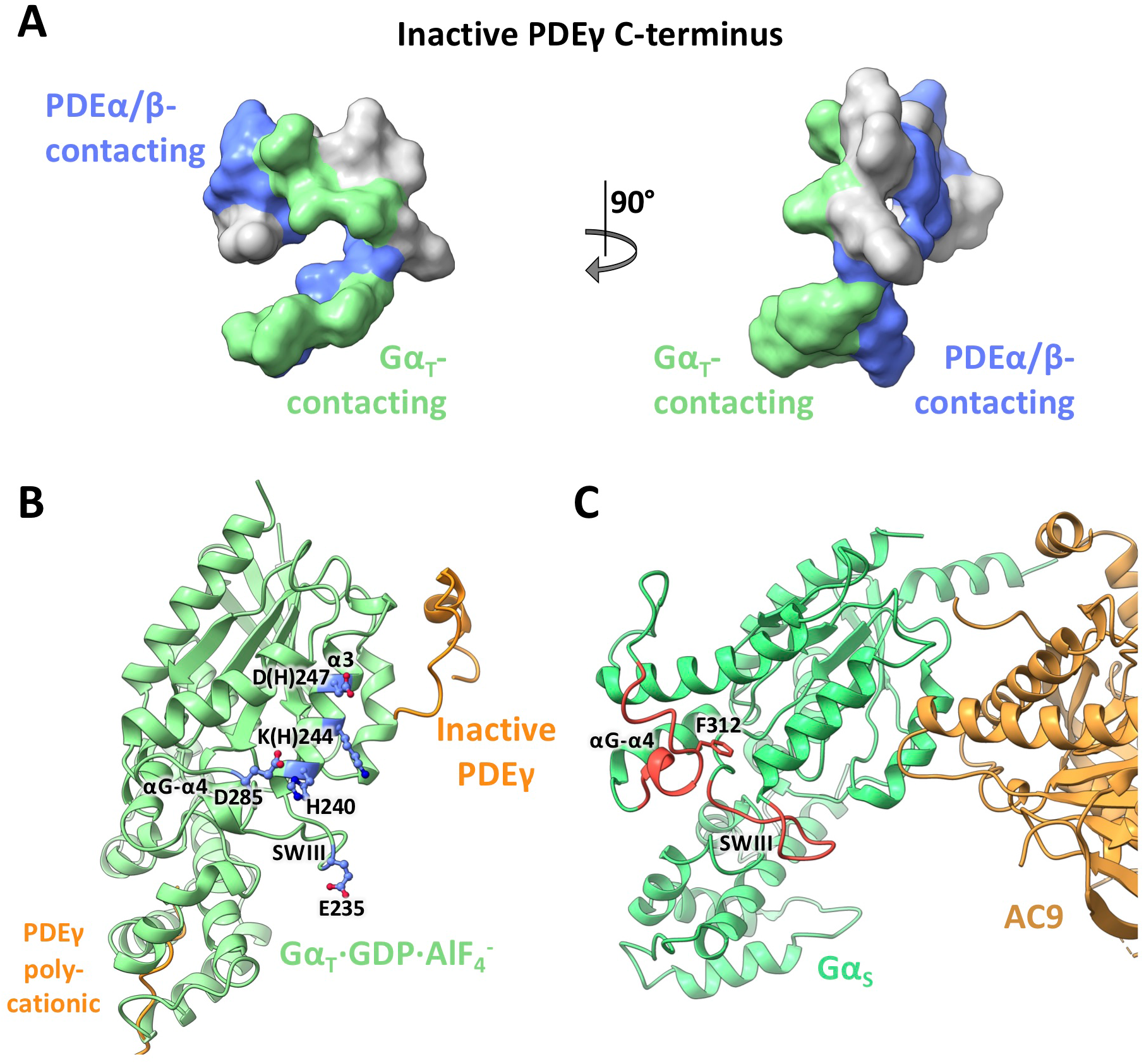
Structural features of the Gα_Τ_·GTP-PDE6 interactions and comparison with the Gα_S_-AC9 complex, Related to Figure 3. (**A**) Interactions between the PDEγ C-terminus and PDEα/β or Gα_Τ_·GTP. The PDEγ C-terminus peptide is shown as grey density, with its PDEα/β-interacting surface colored blue according to the inactive PDE6 structure (PDB: 6MZB) and its Gα_Τ_·GTP-interacting surface colored green based on the Gα_T_·GTP-PDE complex structure. (**B**) The Gα_T_·GDP·AlF_4_^−^ structure from the Gα_T_·GDP·AlF_4_^−^-PDEγ C-ter-RGS9 domain complex (PDB: 1FQJ) aligned onto the inactive PDEγ structure (PDB:6MZB) based on PDEγ C-ter. Residues shown interacting with the PDEγ polycationic region (colored blue) in the Gα_T_·GTP-PDE complex structure (Figure 3C) are located far away from this region in this conformation. (**C**) Position of the homologous regions (colored red) in Gα_S_ in the Gα_S_-AC9 complex corresponding to those in Gα_Τ_·GTP that interact with the PDEγ polycationic region.

**Figure S5.**
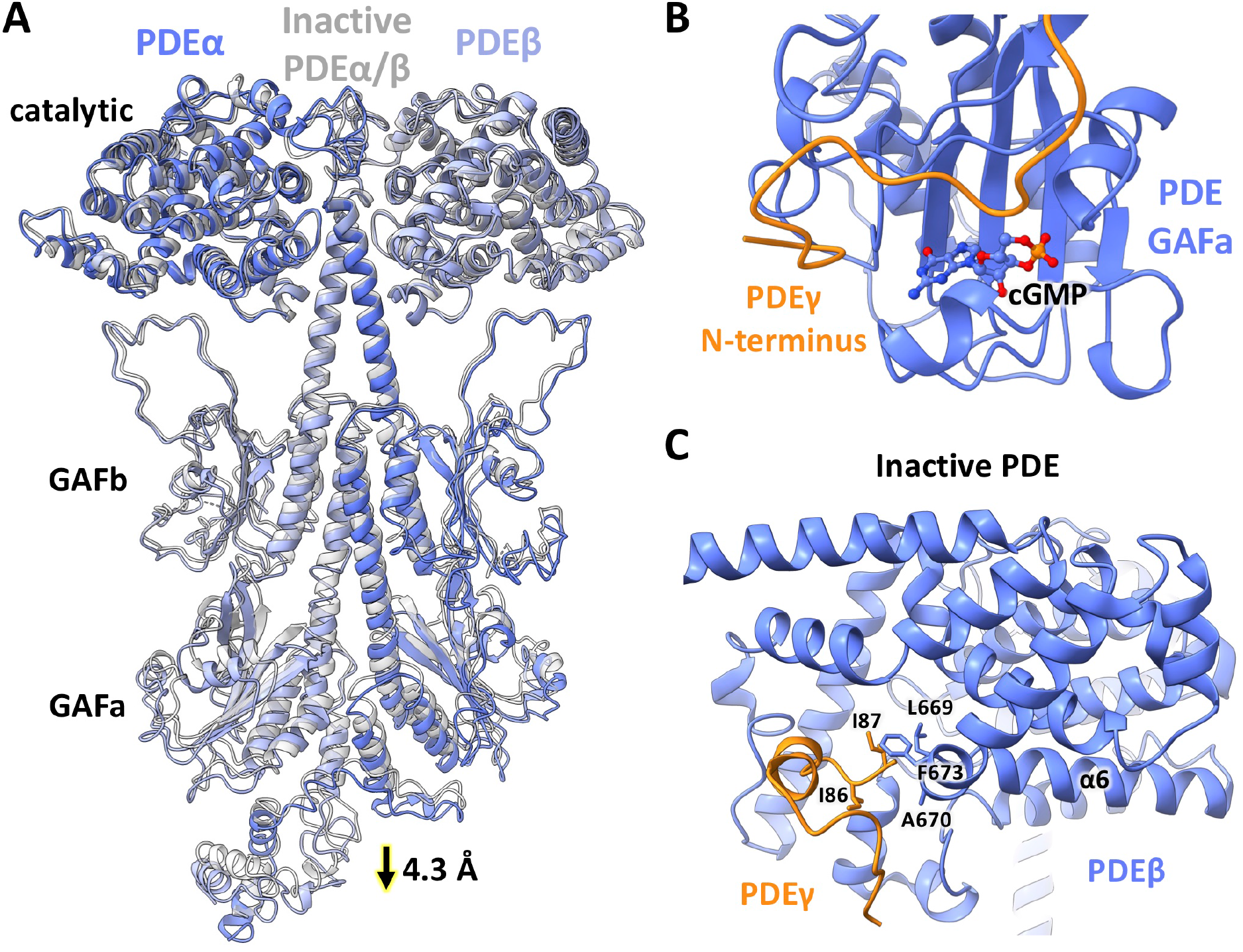
Comparison of the catalytic subunits in the Gα_T_·GTP-PDE6 complex and inactive PDE6, Related to Figures 3 and 5. (**A**) Comparison between the PDEα and PDEβ subunits in the Gα_T_·GTP-PDE6 complex and inactive PDE6 (PDB:6MZB) based on alignment of the catalytic domains. (**B**) The GAFa cGMP-binding pocket in the Gα_T_·GTP-PDE complex. (**C**) Interaction between the α6 helix of PDEα/β and the PDEγ C-terminus in the inactive PDE6 structure (PDB:6MZB).

**Figure S6.**
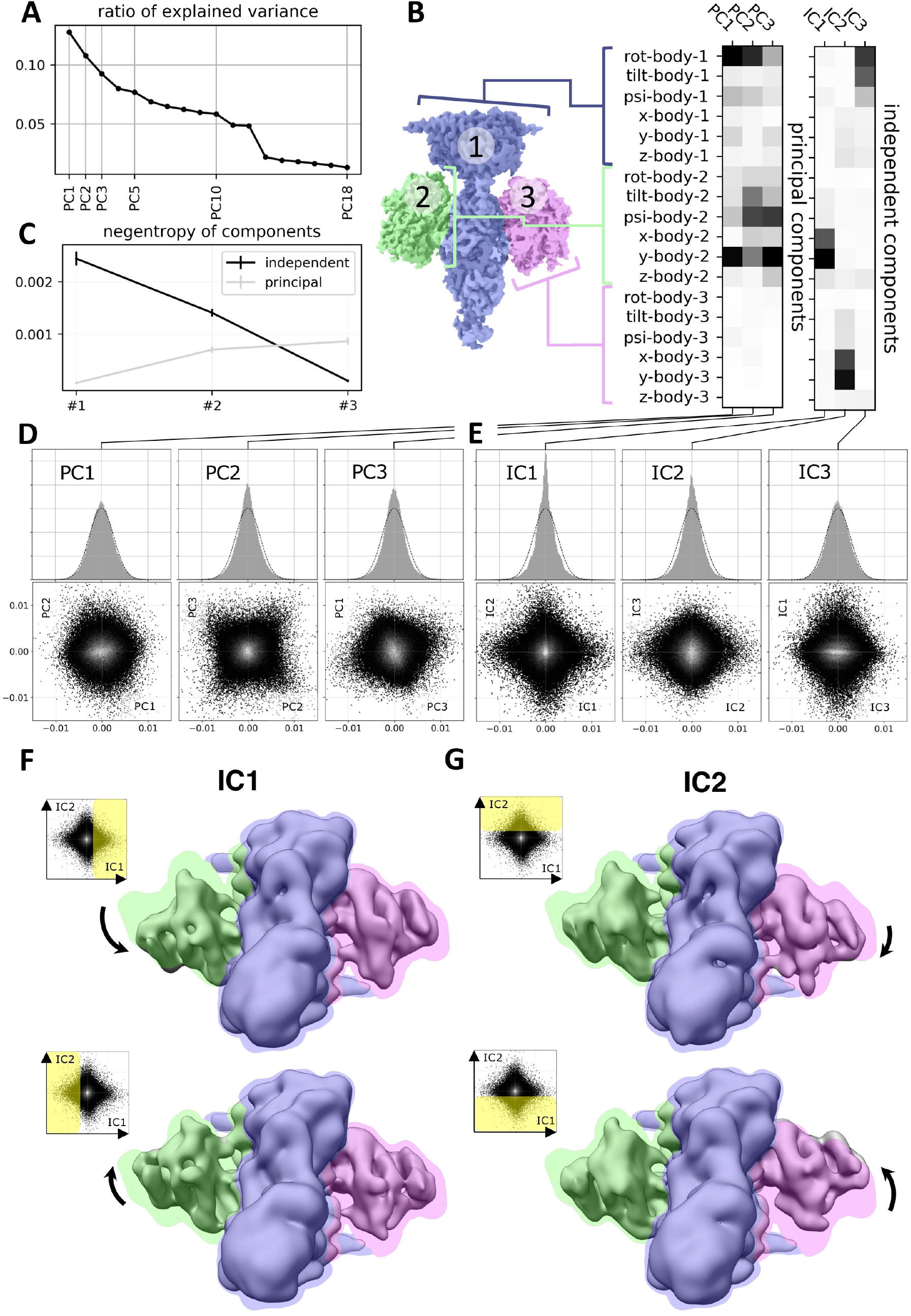
Flexibility analysis results, Related to Figures 5 and 6. (A) Principal component analysis of the 18 multi-body parameters refined for each particle image yields 18 principal components (PC) displayed here in decreasing order of explained variance. The first 3 components explain more than 30% of the variability in the particle images. (B) (left) definition of the multi-body segmentation: the central PDE6 stalk in blue corresponds to Body 1, while the two Gα_T_·GTP subunits correspond to Bodies 2 and 3. (right) The motion of each body is parameterized with 3 translational parameters and 3 rotational parameters. Each of the 18 principal and 3 independent components is a linear combination of the resulting 18 rigid-body parameters, and their weights are shown here for the first 3 principal components (from negligible to larger weight as the shade of grey becomes darker). (C) Negentropy (i.e. reverse entropy) of the first 3 principal and independent components. (D) (resp. (E)) – (top) histogram of the projection of all image particle parameters on the first 3 principal (resp. independent) components PC1, PC2 and PC3 (resp. IC1, IC2 and IC3). (bottom) 2D histograms of the projections of all image particle parameters on all pairs of the first 3 principal (resp. independent) components. (F) (resp. (G)) – Maps illustrating the motions carried by IC1 (resp. IC2). (top) map reconstructed from the particles whose projections belong to the last bin along IC1 (resp. IC2). (bottom) map reconstructed from the particles whose projections belong to the first bin along IC1 (resp. IC2). All maps are shown overlaid on the consensus map, with threshold set at a lower density value, colored according to the scheme in B.

**Figure S7.**
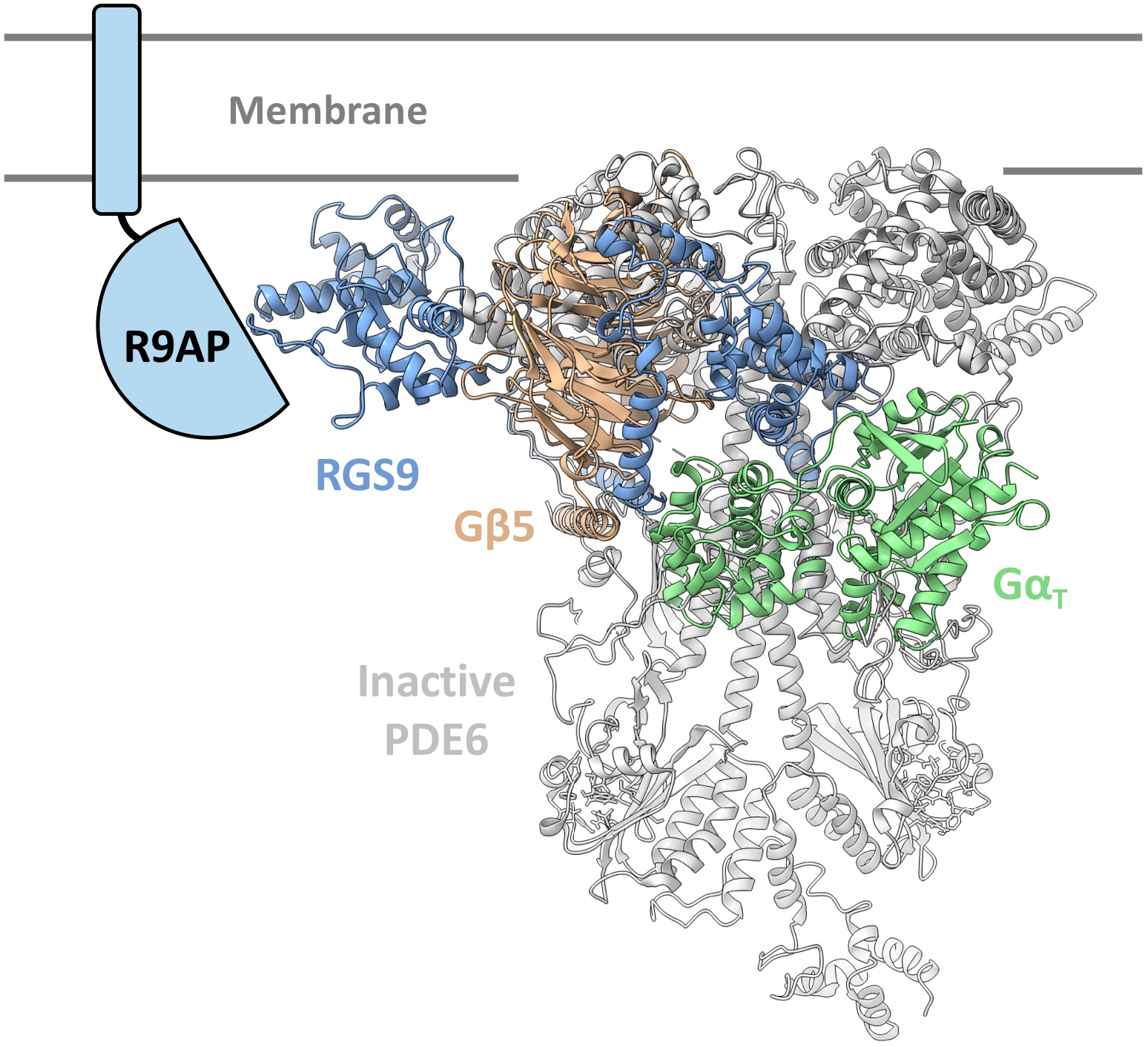
Structural model aligning Gα_T_ and the RGS9 complex onto inactive PDE6, Related to Figures 5 and 6. The Gα_T_·GDP·AlF_4_^−^ structure (colored green) from the Gα_T_·GDP·AlF_4_^−^-PDEγ C-ter-RGS9 domain complex (PDB: 1FQJ) is superimposed onto the inactive PDE structure (colored gray, PDB:6MZB), based on alignment of the PDEγ C-ter peptide. The RGS9/Gβ5 complex structure (RGS9 colored in blue, Gβ5 colored in beige; PDB: 2PBI) is aligned onto the Gα_T_·GDP·AlF_4_^−^ structure based on the C-terminal RGS9 domain. R9AP, which binds at the N-terminal DEP-DHEX domains of RGS9, is shown as a cartoon illustration, colored in cyan.

**Table S1.** CryoEM data collection, refinement and validation statistics.

| | $G\alpha_T$-GTP-PDE6 Complex |
| --- | --- |
| **Data collection and processing** |  |
| Magnification | 29,000 |
| Voltage (kV) | 300 |
| Electron exposure ($e^-/\text{\AA}^2$) | 48 |
| Defocus range ($\mu\text{m}$) | -1.5~-2.5 |
| Pixel size ($\text{\AA}$) | 0.8521 |
| Symmetry imposed | C1 |
| Initial particle images (no.) | 841,967 |
| Final particle images (no.) | 143,125 |
| Map resolution ($\text{\AA}$) | 3.2 |
| FSC threshold | (0.143) |
| **Refinement** |  |
| Initial model used (PDB code) | 6MZB, 1TND |
| Model composition |  |
| Non-hydrogen atoms | 19659 |
| Protein residues | 2404 |
| Ligands | 10 |
| B factors ($\text{\AA}^2$) | |
| Protein | 84.5 |
| Ligand | 85.1 |
| R.m.s. deviations |  |
| Bond lengths ($\text{\AA}$) | 0.005 |
| Bond angles ($^\circ$) | 0.640 |
| Validation |  |
| MolProbity score | 1.61 |
| Clashscore | 5.92 |
| Poor rotamers (%) | 0.88 |
| EMRinger score | 1.78 |
| Ramachandran plot |  |
| Favored (%) | 95.85 |
| Allowed (%) | 4.11 |
| Outliers (%) | 0.04 |

## ACKNOWLEDGMENTS

This work is supported by the NIH (R35 GM122575 and R01 CA201402 to R.A.C., and R01 NS092695 to G.S.).

## AUTHOR CONTRIBUTIONS

Y.G. developed the purification strategy, performed complex purification and PDE6 activity assays for mutants, built and refined the structural model from the cryoEM map and wrote the first draft of the manuscript. G.E. processed the cryoEM data, obtained the cryoEM map and assisted in the flexibility analysis. S.R. generated the 1D4-tagged transducin construct and performed PDE activity assays with varying transducin concentrations. F.P. performed the flexibility analysis. A.B.S. assisted in cryoEM data processing. O.P. froze grids and obtained cryoEM data. Y.G., G.S. and R.A.C. edited the manuscript with contributions from G.E., S.R. and F.P.. G.S. and R.A.C. supervised the project.

## DATA AND SOFTWARE AVAILABILITY

The cryo-EM density map and atomic model have been deposited in EM Data Bank and Protein Data Bank, respectively.

## REFERENCES

Adams, P.D., Afonine, P.V., Bunkóczi, G., Chen, V.B., Davis, I.W., Echols, N., Headd, J.J., Hung, L.-W., Kapral, G.J., Grosse-Kunstleve, R.W., et al. (2010). PHENIX: a comprehensive Python-based system for macromolecular structure solution. Acta Cryst D, Acta Cryst Sect D, Acta Crystallogr D, Acta Crystallogr Sect D, Acta Crystallogr D Biol Crystallogr, Acta Crystallogr Sect D Biol Crystallogr 66, 213–221.

Artemyev, N.O., and Hamm, H.E. (1992). Two-site high-affinity interaction between inhibitory and catalytic subunits of rod cyclic GMP phosphodiesterase. Biochem J 283, 273–279.

Barad, B.A., Echols, N., Wang, R.Y.-R., Cheng, Y., DiMaio, F., Adams, P.D., and Fraser, J.S. (2015). EMRinger: side chain–directed model and map validation for 3D cryo-electron microscopy. Nature Methods 12, 943–946.

Berlot, C.H., and Bourne, H.R. (1992). Identification of effector-activating residues of Gsα. Cell 68, 911–922.

Cheever, M.L., Snyder, J.T., Gershburg, S., Siderovski, D.P., Harden, T.K., and Sondek, J. (2008). Crystal structure of the multifunctional Gβ5–RGS9 complex. Nat Struct Mol Biol 15, 155–162.

Chen, V.B., Arendall, W.B., Headd, J.J., Keedy, D.A., Immormino, R.M., Kapral, G.J., Murray, L.W., Richardson, J.S., and Richardson, D.C. (2010). MolProbity: all-atom structure validation for macromolecular crystallography. Acta Cryst D 66, 12–21.

Clerc, A., Catty, P., and Bennett, N. (1992). Interaction between cGMP-phosphodiesterase and transducin alpha-subunit in retinal rods. A cross-linking study. J. Biol. Chem. 267, 19948–19953.

Colwell, L.J., Qin, Y., Huntley, M., Manta, A., and Brenner, M.P. (2014). Feynman-Hellmann Theorem and Signal Identification from Sample Covariance Matrices. Phys. Rev. X 4, 031032.

Emsley, P., and Cowtan, K. (2004). Coot: model-building tools for molecular graphics. Acta Crystallogr. D Biol. Crystallogr. 60, 2126–2132.

Gao, Y., Hu, H., Ramachandran, S., Erickson, J.W., Cerione, R.A., and Skiniotis, G. (2019). Structures of the Rhodopsin-Transducin Complex: Insights into G-Protein Activation. Molecular Cell 75, 781–790.e3.

García-Nafría, J., and Tate, C.G. (2019). Cryo-EM structures of GPCRs coupled to Gs, Gi and Go. Molecular and Cellular Endocrinology 488, 1–13.

Gulati, S., Palczewski, K., Engel, A., Stahlberg, H., and Kovacik, L. (2019). Cryo-EM structure of phosphodiesterase 6 reveals insights into the allosteric regulation of type I phosphodiesterases. Science Advances 5, eaav4322.

He, W., Cowan, C.W., and Wensel, T.G. (1998). RGS9, a GTPase Accelerator for Phototransduction. Neuron 20, 95–102.

Hu, G., and Wensel, T.G. (2002). R9AP, a membrane anchor for the photoreceptor GTPase accelerating protein, RGS9-1. PNAS 99, 9755–9760.

Hyvarinen, A. (1999). Fast and robust fixed-point algorithms for independent component analysis. IEEE Transactions on Neural Networks 10, 626–634.

Irwin, M.J., Gupta, R., Gao, X.-Z., Cahill, K.B., Chu, F., and Cote, R.H. (2019). The molecular architecture of photoreceptor phosphodiesterase 6 (PDE6) with activated G protein elucidates the mechanism of visual excitation. J. Biol. Chem. 294, 19486–19497.

Kato, H.E., Zhang, Y., Hu, H., Suomivuori, C.-M., Kadji, F.M.N., Aoki, J., Kumar, K.K., Fonseca, R., Hilger, D., Huang, W., et al. (2019). Conformational transitions of a neurotensin receptor 1–G i1 complex. Nature 572, 80–85.

Krishna Kumar, K., Shalev-Benami, M., Robertson, M.J., Hu, H., Banister, S.D., Hollingsworth, S.A., Latorraca, N.R., Kato, H.E., Hilger, D., Maeda, S., et al. (2019). Structure of a Signaling Cannabinoid Receptor 1-G Protein Complex. Cell 176, 448–458.e12.

Liu, W., and Northup, J.K. (1998). The helical domain of a G protein α subunit is a regulator of its effector. PNAS 95, 12878–12883.

Lyon, A.M., Dutta, S., Boguth, C.A., Skiniotis, G., and Tesmer, J.J.G. (2013). Full-length Gα_q_– phospholipase C-β3 structure reveals interfaces of the C-terminal coiled-coil domain. Nsmb 20, 355–362.

Maeda, S., Qu, Q., Robertson, M.J., Skiniotis, G., and Kobilka, B.K. (2019). Structures of the M1 and M2 muscarinic acetylcholine receptor/G-protein complexes. Science 364, 552–557.

Majumdar, S., Ramachandran, S., and Cerione, R.A. (2006). New Insights into the Role of Conserved, Essential Residues in the GTP Binding/GTP Hydrolytic Cycle of Large G Proteins. J. Biol. Chem. 281, 9219–9226.

Maurice, D.H., Ke, H., Ahmad, F., Wang, Y., Chung, J., and Manganiello, V.C. (2014). Advances in targeting cyclic nucleotide phosphodiesterases. Nat Rev Drug Discov 13, 290–314.

Milano, S.K., Wang, C., Erickson, J.W., Cerione, R.A., and Ramachandran, S. (2018). Gain-of-function screen of α-transducin identifies an essential phenylalanine residue necessary for full effector activation. J. Biol. Chem. 293, 17941–17952.

Molday, L.L., and Molday, R.S. (2014). 1D4 – A Versatile Epitope Tag for the Purification and Characterization of Expressed Membrane and Soluble Proteins. Methods Mol Biol 1177, 1–15.

Moschos, M.M., and Nitoda, E. (2016). Pathophysiology of visual disorders induced by phosphodiesterase inhibitors in the treatment of erectile dysfunction.

Nakane, T., Kimanius, D., Lindahl, E., and Scheres, S.H. (2018). Characterisation of molecular motions in cryo-EM single-particle data by multi-body refinement in RELION. ELife 7, e36861.

Natochin, M., Granovsky, A.E., and Artemyev, N.O. (1998). Identification of Effector Residues on Photoreceptor G Protein, Transducin. J. Biol. Chem. 273, 21808–21815.

Neubert, T.A., Johnson, R.S., Hurley, J.B., and Walsh, K.A. (1992). The rod transducin alpha subunit amino terminus is heterogeneously fatty acylated. J. Biol. Chem. 267, 18274–18277.

Noel, J.P., Hamm, H.E., and Sigler, P.B. (1993). The 2.2 Å crystal structure of transducin-α complexed with GTPγS. Nature 366, 654–663.

Oldham, W.M., and Hamm, H.E. (2008). Heterotrimeric G protein activation by G-protein-coupled receptors. Nature Reviews Molecular Cell Biology 9, 60–71.

Pedregosa, F., Varoquaux, G., Gramfort, A., Michel, V., Thirion, B., Grisel, O., Blondel, M., Prettenhofer, P., Weiss, R., Dubourg, V., et al. Scikit-learn: Machine Learning in Python. MACHINE LEARNING IN PYTHON 6.

Pettersen, E.F., Goddard, T.D., Huang, C.C., Couch, G.S., Greenblatt, D.M., Meng, E.C., and Ferrin, T.E. (2004). UCSF Chimera--a visualization system for exploratory research and analysis. J Comput Chem 25, 1605–1612.

Phillips, W.J., Trukawinski, S., and Cerione, R.A. (1989). An antibody-induced enhancement of the transducin-stimulated cyclic GMP phosphodiesterase activity. J. Biol. Chem. 264, 16679–16688.

Qi, C., Sorrentino, S., Medalia, O., and Korkhov, V.M. (2019). The structure of a membrane adenylyl cyclase bound to an activated stimulatory G protein. Science 364, 389–394.

Qureshi, B.M., Behrmann, E., Schöneberg, J., Loerke, J., Bürger, J., Mielke, T., Giesebrecht, J., Noé, F., Lamb, T.D., Hofmann, K.P., et al. (2018). It takes two transducins to activate the cGMP-phosphodiesterase 6 in retinal rods. Open Biology 8, 180075.

Rarick, H.M., Artemyev, N.O., and Hamm, H.E. (1992). A site on rod G protein alpha subunit that mediates effector activation. Science 256, 1031–1033.

Simon, M.I., Strathmann, M.P., and Gautam, N. (1991). Diversity of G proteins in signal transduction. Science 252, 802–808.

Skiba, N.P., Bae, H., and Hamm, H.E. (1996). Mapping of Effector Binding Sites of Transducin α-Subunit Using Gαt/Gαil Chimeras. J. Biol. Chem. 271, 413–424.

Slep, K.C., Kercher, M.A., He, W., Cowan, C.W., Wensel, T.G., and Sigler, P.B. (2001). Structural determinants for regulation of phosphodiesterase by a G protein at 2.0 Å. Nature 409, 1071.

Stryer, L. (1991). Visual excitation and recovery. J. Biol. Chem. 266, 10711–10714.

Sung, B.-J., Hwang, K.Y., Jeon, Y.H., Lee, J.I., Heo, Y.-S., Kim, J.H., Moon, J., Yoon, J.M., Hyun, Y.-L., Kim, E., et al. (2003). Structure of the catalytic domain of human phosphodiesterase 5 with bound drug molecules. Nature 425, 98–102.

Tejada, I.S. de, Angulo, J., Cuevas, P., Fernández, A., Moncada, I., Allona, A., Lledó, E., Körschen, H.G., Niewöhner, U., Haning, H., et al. (2001). The phosphodiesterase inhibitory selectivity and the in vitro and in vivo potency of the new PDE5 inhibitor vardenafil. Int J Impot Res 13, 282–290.

van der Walt, S., Colbert, S.C., and Varoquaux, G. (2011). The NumPy Array: A Structure for Efficient Numerical Computation. Computing in Science Engineering 13, 22–30.

Wensel, T.G., and Stryer, L. (1986). Reciprocal control of retinal rod cyclic GMP phosphodiesterase by its γ subunit and transducin. Proteins: Structure, Function, and Bioinformatics 1, 90–99.

Zhang, K. (2016). Gctf: Real-time CTF determination and correction. Journal of Structural Biology 193, 1–12.

Zhang, X.-J., Cahill, K.B., Elfenbein, A., Arshavsky, V.Y., and Cote, R.H. (2008). Direct Allosteric Regulation between the GAF Domain and Catalytic Domain of Photoreceptor Phosphodiesterase PDE6. J. Biol. Chem. 283, 29699–29705.

Zheng, S.Q., Palovcak, E., Armache, J.-P., Verba, K.A., Cheng, Y., and Agard, D.A. (2017). MotionCor2: anisotropic correction of beam-induced motion for improved cryo-electron microscopy. Nature Methods 14, 331–332.

Zivanov, J., Nakane, T., Forsberg, B.O., Kimanius, D., Hagen, W.J., Lindahl, E., and Scheres, S.H. (2018). New tools for automated high-resolution cryo-EM structure determination in RELION-3. ELife 7, e42166.

